# Protein-protein interactions with fructose-1-kinase alter function of the central *Escherichia coli* transcription regulator, Cra

**DOI:** 10.1101/201277

**Authors:** Dipika Singh, Max S. Fairlamb, Kelly S. Harrison, Chamitha Weeramange, Sarah Meinhardt, Sudheer Tungtur, Benjamin F. Rau, P. Scott Hefty, Aron W. Fenton, Liskin Swint-Kruse

## Abstract

In *E. coli*, the master transcription regulator Cra regulates >100 genes in central metabolism by binding upstream DNA operator sequences. Genes encoding glycolytic enzymes are repressed, whereas those for gluconeogenesis and the citric acid cycle are activated. Cra-DNA binding is allosterically diminished by binding to either fructose-1-phosphate (F-1-P, generated upon fructose import) or fructose-1,6-bisphosphate (F-1,6-BP). F-1,6-BP is generated from F-1-P by the enzyme fructose-1-kinase (FruK) or from other sugars and is a key intermediate in glycolysis. Here, we report that Cra directly interacts with FruK to form a tight protein-protein complex. Further, growth assays with a *fruK* knockout strain show that FruK has a broader role in metabolism than its known role in fructose catabolism. Biochemical experiments show that F-1,6-BP binding enhances either the Cra/FruK interaction and/or CRA binding to DNA and that FruK can catalyze the reverse reaction of F-1,6-BP to F-1-P. Results were used to propose a model in which the Cra-FruK complex enhances activation of gluconeogenic genes. Finally, since FruK itself is repressed by Cra, these newly-reported events add layers to the dynamic regulation of *E. coli* central metabolism that occur in response to changing nutrients.

---

In *Escherichia coli*, more than 100 genes encoding the enzymes of central metabolism are directly regulated by the <u>c</u>atabolite <u>r</u>epressor <u>a</u>ctivator protein (“Cra”). Cra functions as either an activator or a repressor of transcription, depending upon the juxtaposition of the *cra* operators and the gene promoters (Ravcheev *et al.*, 2014a, Shimada *et al.*, 2011). In general, genes encoding glycolytic enzymes are repressed by Cra, whereas those encoding enzymes for gluconeogenesis and the citric acid cycle are activated (Saier & Ramseier, 1996, Ramseier *et al.*, 1995).

Cra’s effects on transcription are modulated by the presence of fructose metabolites that allosterically diminish DNA binding (Ramseier *et al.*, 1993): The strongest inducer is fructose-1-phosphate (F-1-P), which binds to the Cra regulatory domains with micromolar affinities and greatly diminishes Cra DNA binding (Ramseier *et al.*, 1993). F-1-P is created when fructose is transported into the cell *via* the FruBA phosphotransferase system (Figure 1) (Kornberg, 2001) and is further metabolized by fructose-1-kinase (FruK) to fructose-1,6-bisphophate (F-1,6-BP) (Veiga-da-Cunha *et al.*, 2000, Ferenci & Kornberg, 1973). Millimolar concentrations of F-1,6-BP also induce Cra, although DNA binding is not fully diminished (as compared to induction by F-1-P) (Ramseier *et al.*, 1993). In addition to its formation from F-1-P, F-1,6-BP is generated from several other sugars and is a key intermediate in glycolysis. *In vivo* F-1,6-BP concentrations can reach 15 mM when *E. coli* are grown on glucose (Bennett *et al.*, 2009) (Table 1).

**Figure 1.**
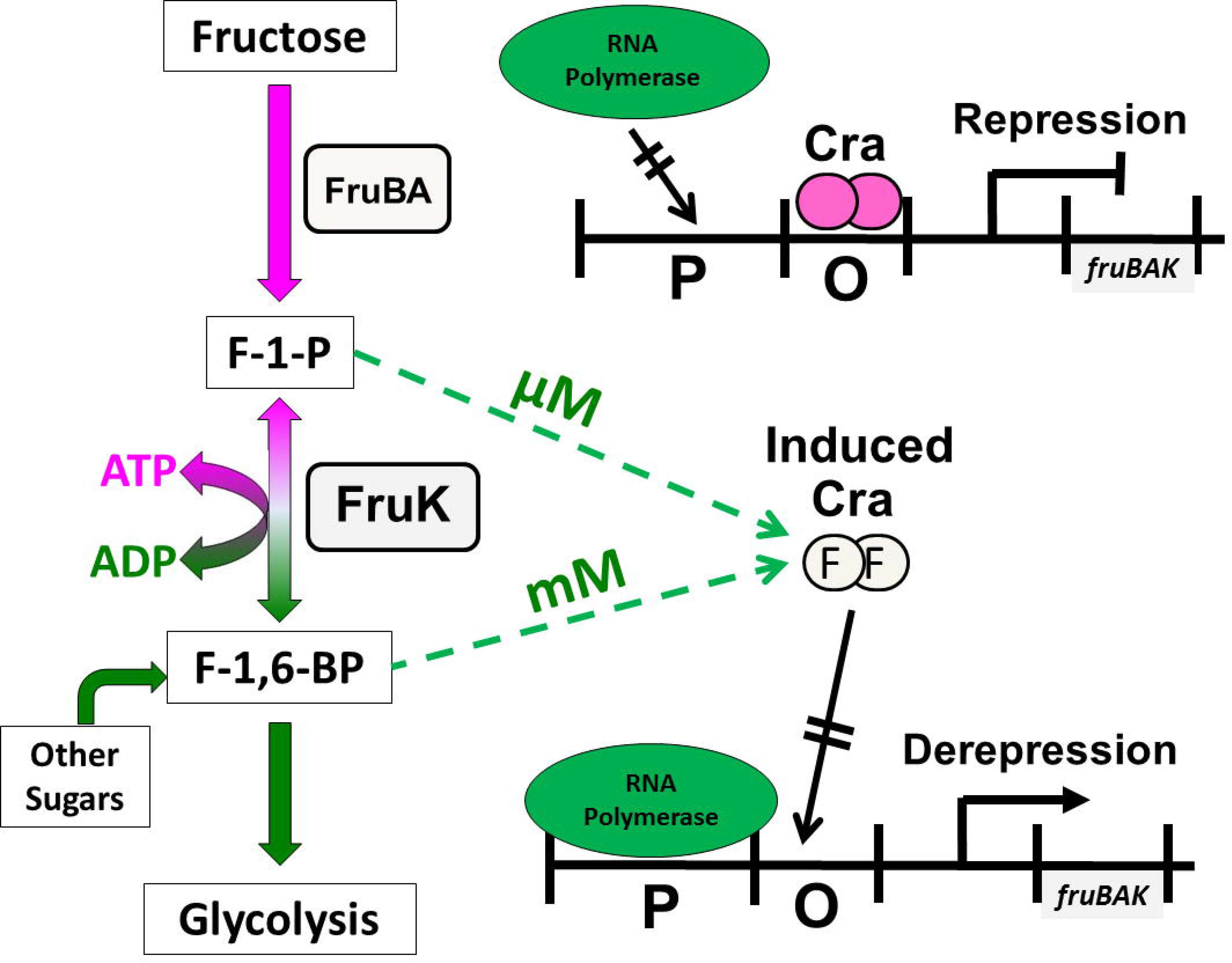
Co-regulation of Cra and FruK. The upper right portion of this cartoon depicts Cra binding to the operator of the *fruBAK* operon to repress downstream genes. When bound to F-1-P at micromolar concentrations or F-1,6-BP at millimolar concentrations, Cra no longer tightly binds the operator and *fruBAK* genes are transcribed (lower right schematic). The gene product of *fruK*, FruK, interconverts the two fructose metabolites.

**Table 1.** Physiological ranges for FruK substrates in *E. coli*^a^

|  | Carbon source (4 g/l in minimal media) |  |  |
| --- | --- | --- | --- |
|  | glucose | glycerol | acetate |
| ATP | 9.6 mM<sup>b</sup> | 9.0 mM<sup>b</sup> | 4.1 mM |
| ADP<sup>c</sup> | 0.55 mM | 0.15 mM | 0.19 mM |
| F-1,6-BP | 15 mM | 5.9 mM | < 0.1 mM |

<sup>a</sup>Values taken from Bennett *et al.* (Bennett *et al.*, 2009). The concentration of F-1-P was not determined in this study.
<sup>b</sup>ATP differences between glucose and glycerol growth conditions were not statistically significant.
<sup>c</sup>ADP differences between the three growth conditions were not statistically significant.

As described above, Cra regulation of gene transcription follows the simple regulatory cycle that is common to homologs of the LacI/GalR family (Swint-Kruse & Matthews, 2009). However, these biochemical features do not fully explain several biological observations for *E.coli.* For example, F-1,6-BP induction of Cra has been used to explain how enterohemorrhagic *E. coli* (EHEC) respond to changes in environmental nutrients (Curtis *et al.*, 2014), but this rationale did not take into account the fact that millimolar F-1,6-BP does not fully induce Cra-DNA binding (Ramseier *et al.*, 1993). In a second example, Cra was proposed to act as a flux sensor that uses intracellular F-1,6-BP concentrations to integrate information from a variety of nutrients (Kochanowski *et al.*, 2013). However, most of the *in vivo* F-1,6-BP levels measured in this study were less than 1 mM, nearly an order of magnitude lower than needed for this metabolite to directly induce Cra (Ramseier *et al.*, 1993). In a third example, a model of *E.coli* metabolic flux used the strong F-1-P inducer as the only Cra modulator, but the model was insufficient to reconcile computations with experiment (Usuda *et al.*, 2010).

Indeed, we find it enigmatic that the strong inducer for Cra (F-1-P) arises only under the rare condition of available fructose, whereas the weaker inducer (F-1,6-BP) is encountered in the presence of many sugars. One possibility is that Cra regulation of metabolism involves other factors. In support of this hypothesis, Cra was shown to directly interact with KdpE for coregulation of a pathogenicity island in EHEC (Njoroge *et al.*, 2012). Here, we report a newly-discovered, high affinity protein-protein interaction between DNA-bound Cra and FruK in nonpathogenic *E. coli* that is enhanced in the presence of F-1,6-BP. Second, we show that deleting the gene for FruK impacts a Cra-associated switch between carbon sources. Together, these findings suggest a model in which FruK and F-1,6-BP binding enhance Cra activation of gluconeogenic pathways. In addition, results provide initial evidence that FruK can catalyze the reverse reaction, converting F-1,6-BP to F-1-P, which provides a potential avenue for forming the strong inducer from a wide range of sugars.

## Results

### Discovery of the Cra/FruK interaction

Discovery of the Cra-FruK interaction arose from our efforts to engineer synthetic LacI/GalR transcription regulators (Shis *et al.*, 2014, Meinhardt *et al.*, 2012). Homologs in the LacI/GalR family have an N-terminal DNA binding domain and a C-terminal regulatory domain that binds allosteric ligands (Swint-Kruse & Matthews, 2009). One of the engineered constructs was created by fusing the LacI DNA binding domain to the Cra regulatory domain to create “LLhF”; this chimeric protein bound *lac* operator sequences and was induced by fructose metabolites (Meinhardt *et al.*, 2012). To verify the *in vivo* activity of LLhF and other LacI/GalR chimeras, the synthetic repressors were expressed in *E. coli* grown in MOPS-glycerol media, and pull-down assays (using *lac* operators immobilized to magnetic beads) were carried out in crude cell extracts (Meinhardt *et al.*, 2012). As expected, wild-type LacI and other chimeras were detected as a single band as compared to control cells (Meinhardt *et al.*, 2012). However, the Cra chimera LLhF showed two bands, with the second band slightly smaller than that for repressor (Figure 2, lane 1). Mass spectrometry was used to identify the second band as fructose-1-kinase (FruK), a member of ribokinase family (Orchard & Kornberg, 1990).

**Figure 2.**
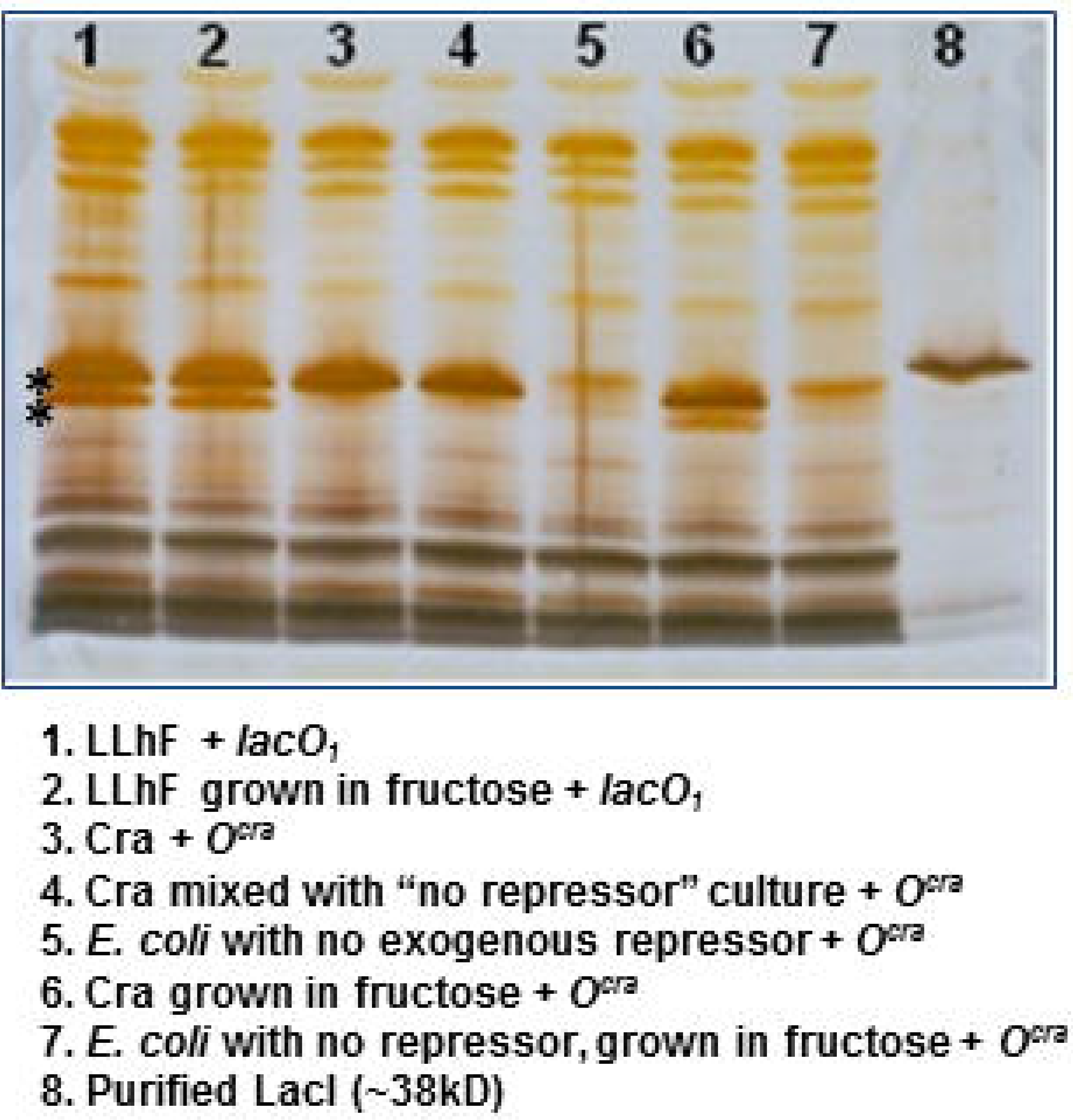
LLhF and Cra form complexes with FruK in *E.coli* crude cell extracts. Either the LLhF chimera or the wild-type Cra was over-expressed in *E.coli* 3.300 cells. Cells were grown in minimal media with glycerol as the carbon source, in either the absence or presence of additional fructose. Lysed cells were incubated with immobilized DNA and the DNA binding proteins were visualized with SDS-PAGE. The asterisks indicate the positions of the repressor and FruK bands. In this experiment, all FruK was expressed from the genomic *fruBAK* operon. In lane 8, purified *E. coli* lactose repressor protein serves as a molecular weight marker; this protein (360 a.a.; 38590 Da monomer; Uniprot P03023) is slightly larger than Cra (334 a.a.; 37999 Da monomer; Uniprot P0ACP1).(The Uniprot Consortium, 2017)

This result was immediately notable because the two proteins indirectly regulate each other’s functions. First, FruK converts the strong Cra inducer F-1-P to the weak inducer F-1,6-BP via the reaction (Ferenci & Kornberg, 1973):

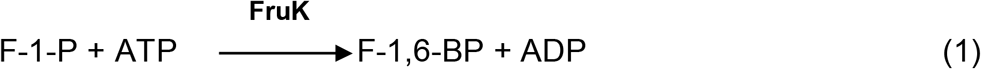

Second, Cra represses the *fruBAK* operon that encodes the FruK protein (Geerse *et al.*, 1989, Geerse *et al.*, 1986, Ramseier *et al.*, 1993). Thus, in addition to the indirect effects, the LLhF-FruK interaction raised the possibility of a direct interaction between wild-type, full-length Cra and FruK.

To test that idea, wild-type Cra was overexpressed in *E.coli* and the pull-down assay was repeated using the consensus, perfectly palindromic (Saier & Ramseier, 1996) *O*^cra^ DNA binding site. In MOPS-glycerol media, cells expressing Cra protein did not pull-down FruK (Figure 2, lane 3). However, when 20 mM fructose was added to the MOPS-glycerol media, the FruK band was again detected as associated with Cra (Figure 2 Lane 6). The different growth conditions required for LLhF and Cra to pull-down FruK lead us to conclude that FruK protein is only detectable when the *fruBAK* operon is de-repressed: For cells over-expressing Cra, derepression was accomplished by the conversion of fructose to inducer F-1-P; in cells overexpressing LLhF, de-repression could be accomplished *via* formation of dominant negative heterodimers between LLhF and endogenous wild-type Cra. (In the LacI/GalR homologs, obligate dimerization occurs via the regulatory domains (Swint-Kruse & Matthews, 2009); mRNA microarray experiments for *E. coli* expressing various LacI/GalR chimeras showed that genes regulated by their respective wild-type homologs were derepressed; this likely occurred through dominant negative heterodimers of the chimeras with endogenous wild-type repressors (unpublished data)).

Since both Cra and the LLhF chimera bound to FruK, we conclude that the kinase binds to the regulatory domain(s) of Cra. To evaluate the binding affinity between wild-type Cra and FruK, pull-down assays were repeated with various concentrations of partially purified Cra and FruK (Supplementary Figure 1). From these experiments, the K_d_ for the interaction was estimated to be <10^−7^ M.

### Deletion of the fruK gene impairs the switch from rich to glycerol minimal media

Based on our discovery of the Cra-FruK complex, we hypothesized that FruK plays a larger role in the Cra regulon. The only widely-known role of FruK has been in fructose metabolism; when fructose is the only available carbon source, deletion/interruption of the *fruK* gene abolishes *E. coli* growth (Ferenci & Kornberg, 1973). If the hypothesis above is correct, then deletion of the *fruK* gene should affect bacterial growth on other carbon sources. Indeed, using a *fruK* mutant strain, Molchanova *et al.* documented a growth defect on lactate and a corresponding decrease in PEP-synthase required for gluconeogenesis (Molchanova *et al.*, 1992). Genes for the latter pathway are activated by Cra (Chin *et al.*, 1989).

To further demonstrate the global metabolic role of FruK, we grew both wild-type and *fruK* deletion *E. coli* strains (see Experimental Procedures for strain genotypes) in LB media and used them to inoculate glucose- and glycerol minimal media. Glucose enters metabolism upstream of F-1,6-BP and proceeds through glycolysis to the TCA cycle. Glycerol enters metabolism downstream of F-1,6-BP, *via* formation of dihydroxyacetone phosphate (DHAP) and can feed into the TCA cycle and can facilitate gluconeogenesis (Figure 3) (Mazumdar *et al.*, 2010, Murarka *et al.*, 2008). Finally, fructose minimal media was used as a control condition.

**Figure 3.**
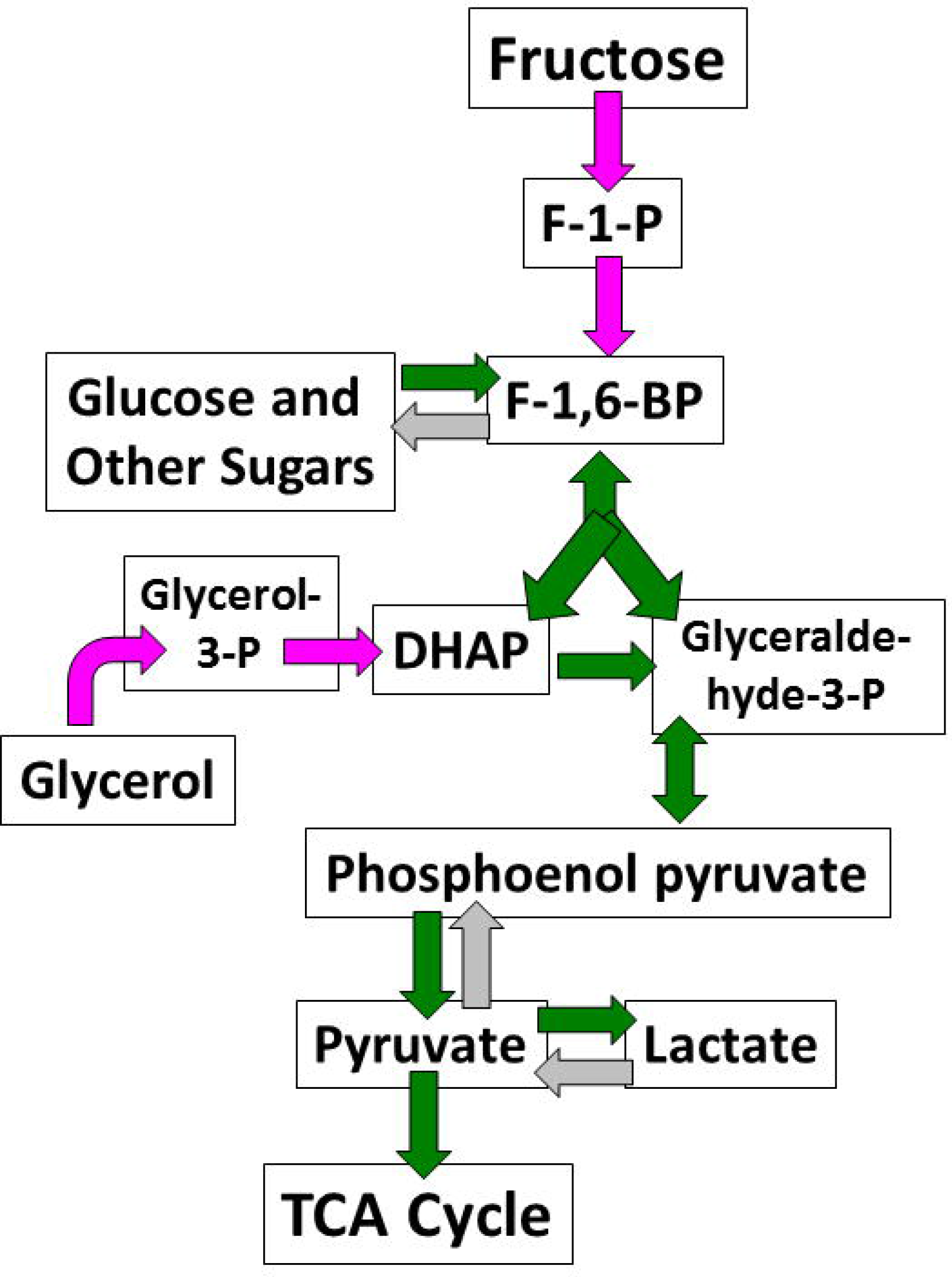
*E. coli* metabolic pathways for glucose, fructose, glycerol, and lactate. Double headed arrows indicate reversible reactions known to be physiologically important. Steps catalyzed specifically for gluconeogenesis are shown with light gray arrows. Information for the figure was derived from references (Mazumdar *et al.*, 2010, Fuchs *et al.*, 2012).

The transition from LB to minimal media resulted in two-phase growth curves (Figure 4) that are reminiscent of the diauxic curves observed when *E. coli* switch between carbon sources (Monod, 1947). In the glucose growth conditions, both the wild-type and *Δfruk* strains showed essentially identical growth curves (Figure 4A). As expected, the *Δfruk* strain showed no growth on fructose after 24 hours, whereas the wild-type strain showed robust growth. However, in glycerol minimal media, the *Δfruk* strain showed a significant increase in the diauxic lag time relative to the wild-type strain (Figure 4B-C). Subsequent to the lag, the rates of the exponential growth phases of the wild-type and deletion strains were very similar to each other (Figure 4D).

**Figure 4.**
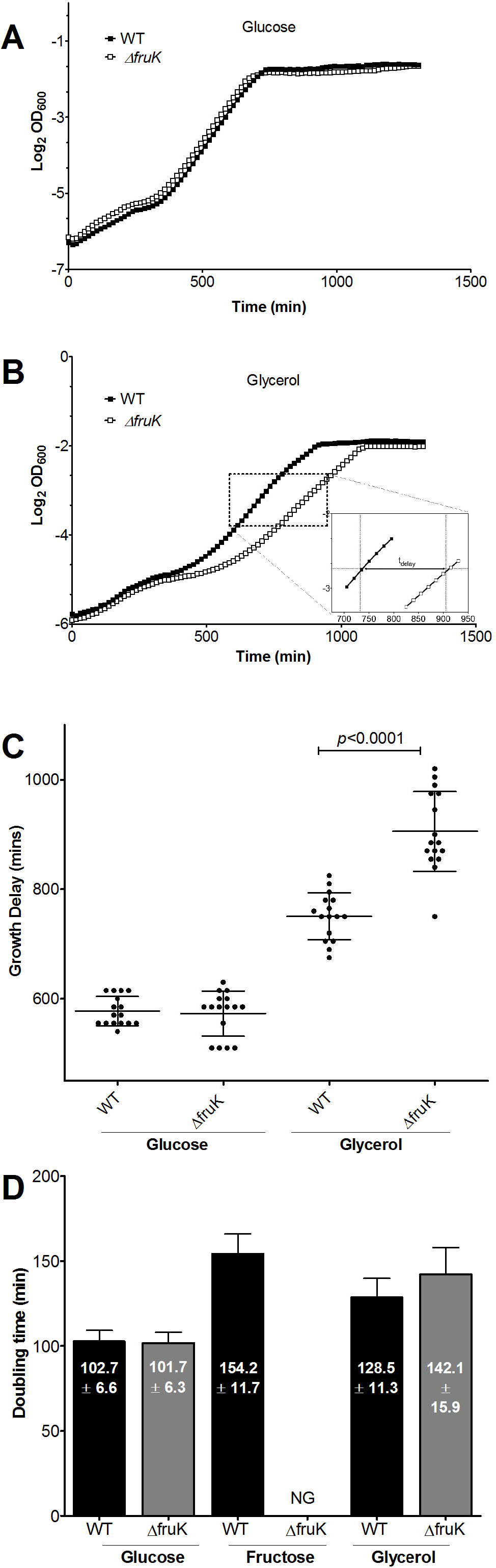
Growth of WT and *ΔfruK* strains on varied carbon sources. WT and *ΔfruK E. coli* were grown in minimal media plus glucose, fructose or glycerol. Plots show the averages of quadruplicate samples for one day’s experiment in (A) glucose and (B) glycerol. Delay time (t_delay_, insert B) was calculated as the amount of time to reach mid-log phase and in (C) is shown as mean and standard deviation of quadruplicate wells from four separate experiments. In glycerol, the mutant showed a significantly increased delay as compared to WT (*p*<0.0001, paired t-test). (D) Doubling times during logarithmic growth are shown as mean and standard deviation of quadruplicate wells from four separate experiments. “NG” indicates “no growth”.

Thus, FruK (most likely *via* the Cra-FruK complex) is involved in the metabolic switch to a carbon source that enters metabolism downstream of F-1,6-BP (Figure 3). To determine possible biochemical roles of Cra-FruK complex formation, we next undertook the complete purification (Supplementary Figure 2) and biochemical characterization of these two proteins and carried out experiments to determine the outcomes of complex formation on DNA binding of Cra and enzyme activity of FruK.

### Confirmation of FruK enzymatic activity

To confirm the kinetic activity of purified FruK, we first replicated the published assay that coupled the FruK reaction to those of aldolase C, glycerophosphate dehydrogenase, and triose phosphate isomerase (Veiga-da-Cunha *et al.*, 2000). From this, we determined a K_m_ value of 0.4 mM, in reasonable agreement with the previously published value of 0.13 mM (Veiga-da-Cunha *et al.*, 2000). However, different lots of commercial muscle aldolase C had varied background activity for F-1-P, which could arise from contamination with other aldolase isozymes (Lebherz & Rutter, 1969). Thus, we established a new assay by coupling the ADP product of the FruK reaction (reaction 1) to the pyruvate kinase (PYK) and lactate dehydrogenase (LDH) reactions and monitoring depletion of NADH at 340 nm (Fenton & Alontaga, 2009).

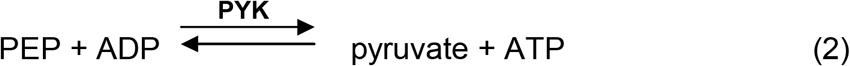

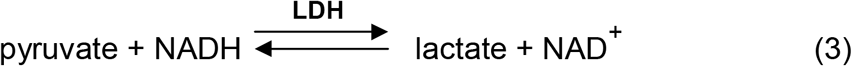

Using several preparations of FruK, we found that K_m_ for F-1-P is in the range of 0.5-2 mM (Figure 5).

**Figure 5.**
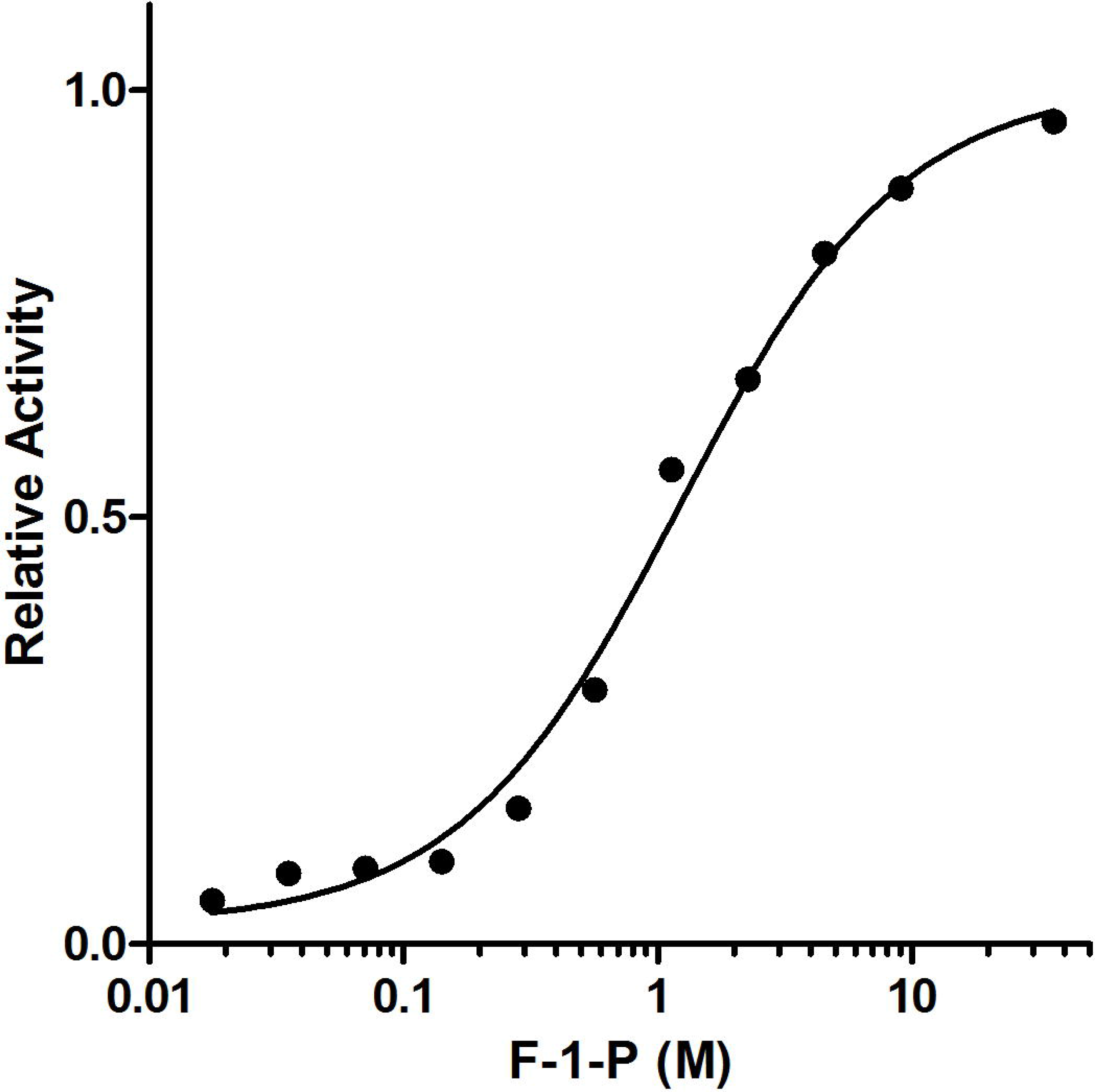
Enzymatic activity of FruK. The catalytic activity of FruK was determined as a function of F-1-P concentration as described in the text. Solid circles represent the measured, initial rates of catalysis, normalized to the maximal value after fitting to the Michaelis-Menton equation. The solid line is the best fit of the Michaelis-Menten equation to the data. Similar responses were seen for several preparations of purified FruK and for FruK that co-purified with wild-type Cra and with the chimera LLhF.

To determine whether the interaction with Cra could regulate the enzymatic activity of FruK, we desired to add purified Cra to purified FruK. However, control reactions showed that Cra was contaminated with small amounts of FruK. The FruK contaminant was confirmed by SDS-PAGE using either over-loaded purified Cra or Cra pull-down samples (*i.e.* Supplementary Figure 1, left gel, lanes 1 and 2). To date, we have been unable to purify Cra from *fruk* deletion strains of *E.coli.* (Although Cra is expressed, it does not survive ammonium sulfate precipitation). Thus, we instead compared K_m_ for F-1-P of purified FruK with that for co-purifying Cra/FruK; values were statistically identical, even in the presence of operator DNA sequences. We were unable to compare values for V_max_ because the extremely low concentration of FruK could not be accurately quantified in the co-purified Cra/FruK sample.

The FruK activity in the co-purified Cra/FruK sample confirms the identity of the FruK protein, and the ever-present contamination of FruK in our Cra preparations provide further evidence to the strength of the Cra/FruK interaction. In addition, results for the FruK activity in the complex have three possible interpretations: (i) The Cra/FruK interaction has no effect on K_m_ for F-1-P. (ii) The Cra/FruK complex cannot form in the presence of high concentrations of F-1-P needed to carry out the assay. (iii) The FruK enzyme activity is only altered when the Cra/FruK complex is bound to DNA; however, the ternary Cra/FruK:DNA complex formation is greatly diminished in the presence of F-1-P (see below) and thus that complex is not likely to assemble in the current enzymatic assay.

### The Cra/FruK binds operator DNA in an F-1,6-BP-dependent manner

To determine the binding affinities (K_d_ values) of purified Cra for operator DNA, equilibrium DNA binding assays were performed. Experiments employed two different techniques: filter binding (Falcon & Matthews, 2001, Riggs *et al.*, 1968, Wong & Lohman, 1993), and biolayer interferometry (Concepcion *et al.*, 2009). Experiments were carried out using operator DNA sequences from three different operons: (i) The *fruB-prox* and *fruB-distal* operators are bound by Cra to repress the *fruBAK* genes (Ramseier *et al.*, 1993); we report herein that both are high affinity operators. (ii) The moderate affinity DNA operator *aceB* is bound by Cra to activate the transcription of malate synthase (Cortay *et al.*, 1994). And (iii) the DNA operator *pfkA* is bound by Cra to repress expression of fructose-6-phosphate kinase; this DNA operator lacks the canonical GC basepair between the inverted repeats (Negre *et al.*, 1996); the K_d_ estimated for *pfkA* is 3.5 nM (Negre *et al.*, 1996).

All experiments were carried out at very low Cra concentrations (50 nM or less); as such, the concentration of co-purifying FruK should be very low. Since adding separately-purified FruK to Cra elicited a large response in biolayer interferometry assays (see below), this assumption appears to be valid.

Filter binding assays under stoichiometric binding conditions (Swint-Kruse & Matthews, 2004) showed that purified Cra comprised 70-90% active repressor protein. For Cra binding to the DNA operator *aceB*, the filter binding technique was used to determine a DNA binding affinity of 1.7 ± 0.5 х 10^−9^ M (Figure 6A), which is in very good agreement with the published value of 3 nM (Ramseier *et al.*, 1993). Filter binding was also used to determine a K_d_ value of 1.8 ± 0.1х10^−11^ M for Cra binding to both the *fruB* distal and proximal DNA operators (Figure 6B).

**Figure 6.**
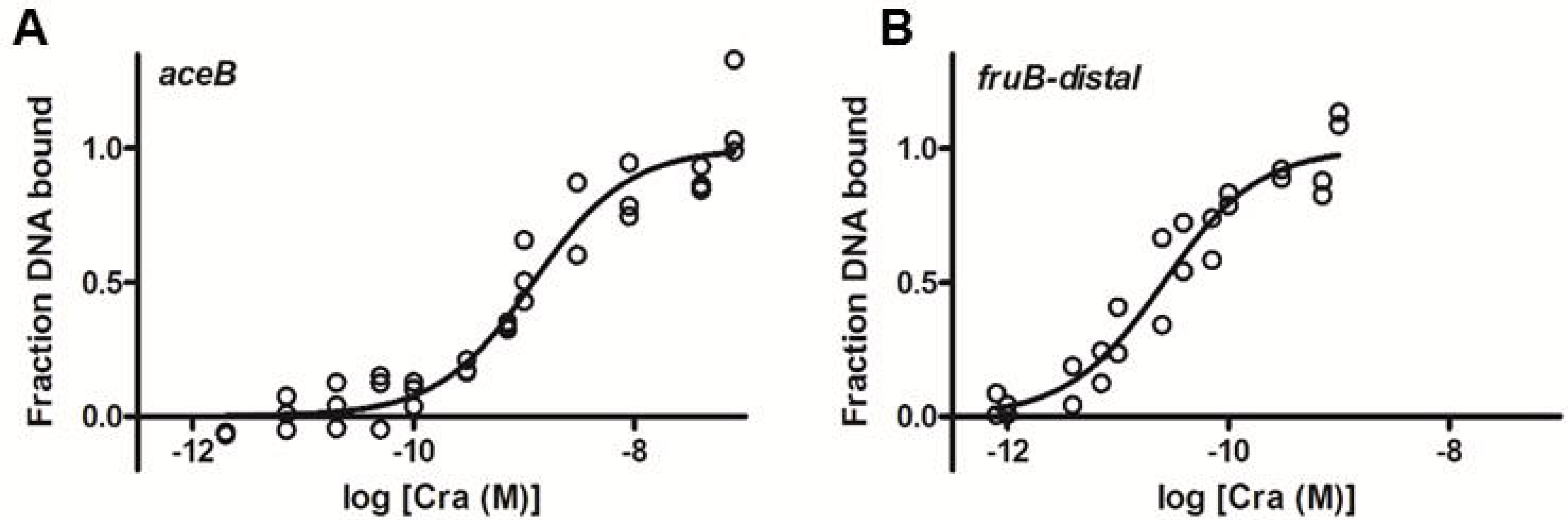
Cra-DNA binding monitored with the filter binding assay. The open circles represent the fraction Cra bound to the indicated operator DNA, after normalizing a fit to the scale of zero to 1. The solid line represents the best fit of the equation Y=Y_max_*([Cra]/(K_d_+[Cra]))+c, where the parameters “Y_max_” and “c” were used for normalization of the raw data and, in the normalized data shown, were set to 1 and 0, respectively. Data shown are from a single determination with triplicate samples at each concentration; results from multiple determinations are summarized in the text. The addition of 1 mM fructose-1-phosphate diminished DNA binding to either operator sequence; K_d_ was beyond the limits of the assay (> 10^−7^ M; data not shown).

When purified FruK was added to determine the effect of complex formation on Cra:DNA binding, the filter was saturated with protein, precluding the use of this assay. Thus, we turned to biolayer interferometry (Concepcion *et al.*, 2009). In this assay, one component of the binding reaction is immobilized to a derivatized fiber-optic tip, which is then dipped into a solution of the second component. Binding is monitored as a function of time *via* changes in interference patterns of reflected light. As a general rule, the signal increases as the thickness of the immobilized and bound material increases.

For this project, we immobilized biotinylated operator DNA to streptavidin tips, which were then incubated with repressor protein in the presence and absence of FruK and various small molecules. We first benchmarked the assay using the well-characterized binding of the lactose repressor protein (LacI; *e.g.* (Swint-Kruse *et al.*, 2003, Zhan *et al.*, 2006, Falcon & Matthews, 2001)) and a LacI:PurR chimera (Zhan *et al.*, 2008) to *lac* operators (Supplementary Figures 3-6). These experiments showed that the plateau of the association phase correlated with the percent of repressor bound (Figure 7 and Supplementary Figure 5). Thus, we used this parameter to monitor effects of F-1-P and F-1,6-BP on DNA binding to Cra and to the Cra/FruK complex. Since the association and dissociation rates of the control proteins and Cra showed complicated kinetic phases (Figure 7B and Supplementary Figures 4-5), we did not quantify rate parameters.

**Figure 7.**
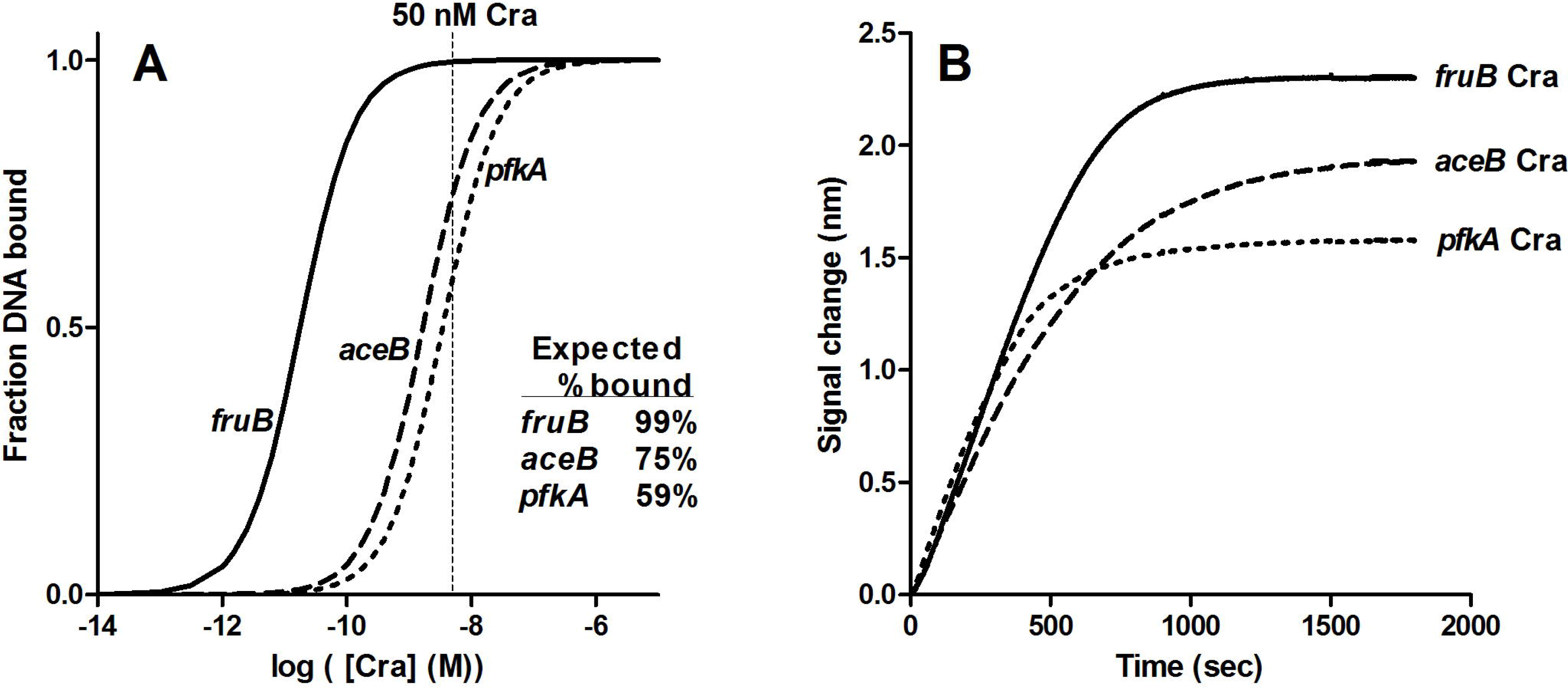
Biolayer interferometry of Cra-DNA binding. (A) Simulated equilibrium binding curves using the Kd values reported in the text for three different Cra operators. The vertical dashed line and the inset table indicate the expected percent bound at the Cra concentration used in biolayer interferometry assay. (B) Biotinylated operator DNA was immobilized to streptavidin tips from an 80 nM solution. DNA binding to 50 nM Cra was monitored with biolayer interferometry. Data were collected every 0.2 seconds and the graph shows the connecting line between data points. The *fruB* sequence used was that of the proximal operator (Table 2). Note that the amplitude at equilibrium correlated with the expected percent bound for the three operators. In control experiments, similar results were observed for the lactose repressor protein binding to its operator DNA sequences (Supplementary Figures 5-6).

Representative data from biolayer interferometry are shown in Figures 7-8 and Supplementary Figure 7. As expected, the interferometry amplitudes at equilibrium correlated with the K_d_ for Cra-DNA binding (Figure 7), and Cra binding to all three operators showed the same response to inducer (Figure 8 and Supplementary Figure 7): When Cra was preincubated with either 1 mM F-1-P or 10 mM F-1,6-BP, the amplitude of the interferometry signal was diminished. This was consistent with the expected behavior of diminished Cra binding to DNA. In parallel with the behavior documented for LacI (Lin & Riggs, 1975a, Lin & Riggs, 1975b, Elf *et al.*, 2007, Taraban *et al.*, 2008) (Supplementary Figure 5), induced Cra retained weak affinity for DNA; the signal did not fall to zero.

**Figure 8.**
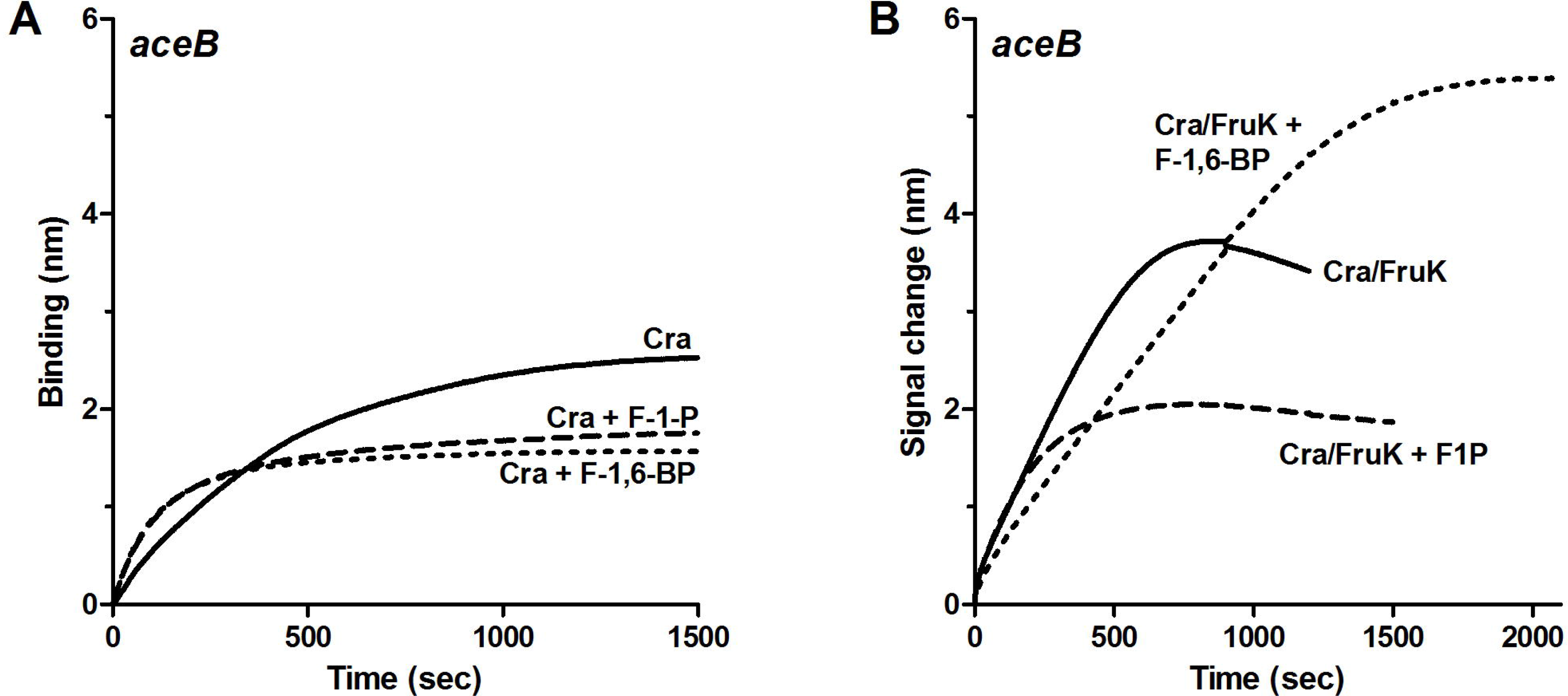
Response of DNA-bound Cra and Cra/FruK complex to fructose metabolites. In both panels, biotinylated *aceB* operator DNA was immobilized to streptavidin tips from an 80 nM solution. (A) 50 nM Cra was bound to the DNA in the absence and presence of pre-equilibrated fructose metabolites. As previously reported for Cra, 1 mM F-1-P and 10 mM F-1,6-BP acted as inducers to diminish DNA binding (dashed and dotted lines). (B) Pre-equilibrated 50 nM Cra and 200 nM FruK was bound to the DNA in the absence and presence of pre-equilibrated fructose metabolites. Note the increased signal of Cra/FruK (solid line) over that for Cra alone (panel A, solid line). The presence of F-1-P appeared to act as an inducer of Cra/FruK, diminishing the amount of complex bound to the tip (dashed line). In contrast, the presence of F-1,6-BP increased the signal (dotted line), which indicates that either more Cra/FruK complex formed, the complex had greater DNA binding, or both.

Next, we pre-incubated Cra with excess FruK and repeated DNA binding experiments. Relative to Cra-only, the amplitude for Cra/FruK binding greatly increased (Figure 8), which is consistent with a larger protein component binding to the immobilized DNA and/or with enhanced DNA binding. (This observation provided evidence that slight contamination of FruK in the purified Cra did not saturate at the Cra concentrations used here.) Control experiments with FruK-only showed no DNA binding, as expected (Supplementary Figure 8).

Pre-incubating Cra/FruK with 1 mM F-1-P or 10 mM F-1,6-BP showed a striking result (Figure 8B): As with Cra only, F-1-P served as an inducer (decreased amplitude). In contrast, the presence of F-1,6-BP increased the amplitude of Cra/FruK binding. This result indicates that F-1,6-BP must either enhance formation of the Cra/FruK complex, enhance DNA binding by the complex, or both. The same trend was observed for all three DNA operator sequences, with two different lengths of flanking DNA. Further, the LLhF chimera showed the same patterns when binding to *lac* DNA operator in the absence and presence of FruK, F-1-P, and F-1,6-BP (Supplementary Figure 9). We note that these experiments do not discriminate whether the effect arises from F-1,6-BP binding to Cra or from binding to FruK.

### The Cra/FruK:DNA complex responds to the forward and reverse reactions of FruK

Since F-1-P and F-1,6-BP have opposite effects on the Cra/FruK:DNA complex, we reasoned that the FruK catalytic reaction could be monitored by biolayer interferometry. To that end we first determined whether the other substrates - ATP and ADP - had effects on Cra or Cra/FruK binding to DNA. Initial experiments in which the protein(s) were pre-incubated with nucleotide showed small variations in the signal amplitude. To better detect any small effects of nucleotide, Cra and Cra/FruK samples were first bound to the immobilized DNA and then ATP (3 mM) or ADP (0.3 mM or 3 mM) was added and the resulting signal change was monitored (Figures 9 and 11). For the Cra:DNA complex, no change was observed from the addition of nucleotide (Figure 9). For the Cra/FruK:DNA complex, addition of 3 mM ATP or ADP caused a slight decrease in the interferometry signal (Figures 9 and 11); addition of 0.3 mM ADP had essentially no impact.

**Figure 9.**
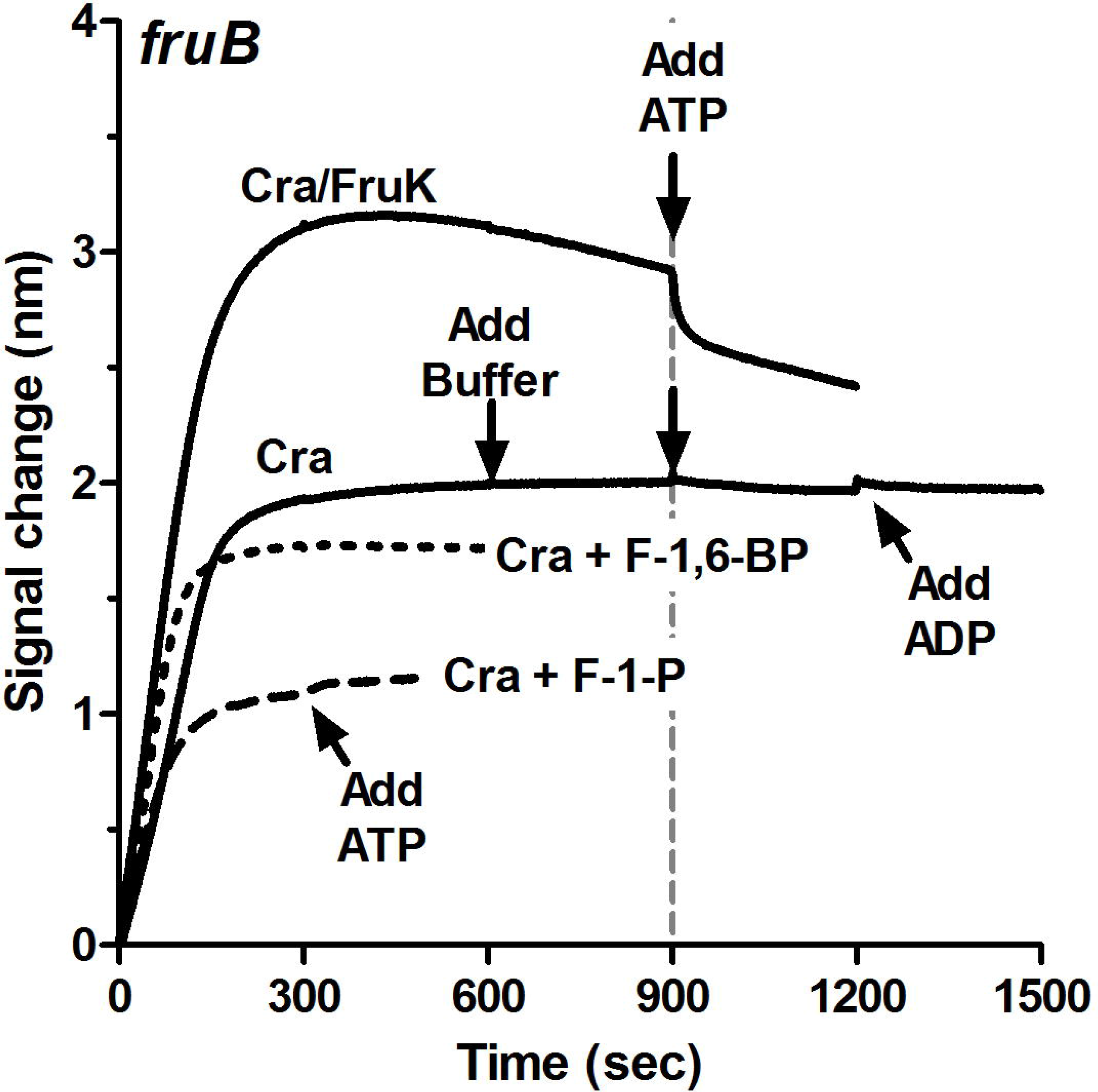
Response of Cra and Cra/FruK complex to ATP and ADP. Biotinylated *fruB-prox* DNA (Table 2) was immobilized to streptavidin tips from an 80 nM solution. Cra (50 nM) or Cra/FruK complex (50 nM/200 nM) was bound to the DNA in the absence and presence of preequilibrated fructose metabolites (1 mM F-1-P or 10 mM F-1,6-BP). At the indicated times, either buffer, 3 mM ATP (dashed gray vertical line), or 3 mM ADP was added to the samples.

Next, we carried out the forward reaction by alternating additions of F-1-P and ATP to the DNA-immobilized Cra/FruK complex (Figure 10). The resulting signal changes were consistent with the forward reaction, in which F-1-P was converted to F-1,6-BP, thereby enhancing the signal for the Cra/FruK:DNA complex.

**Figure 10.**
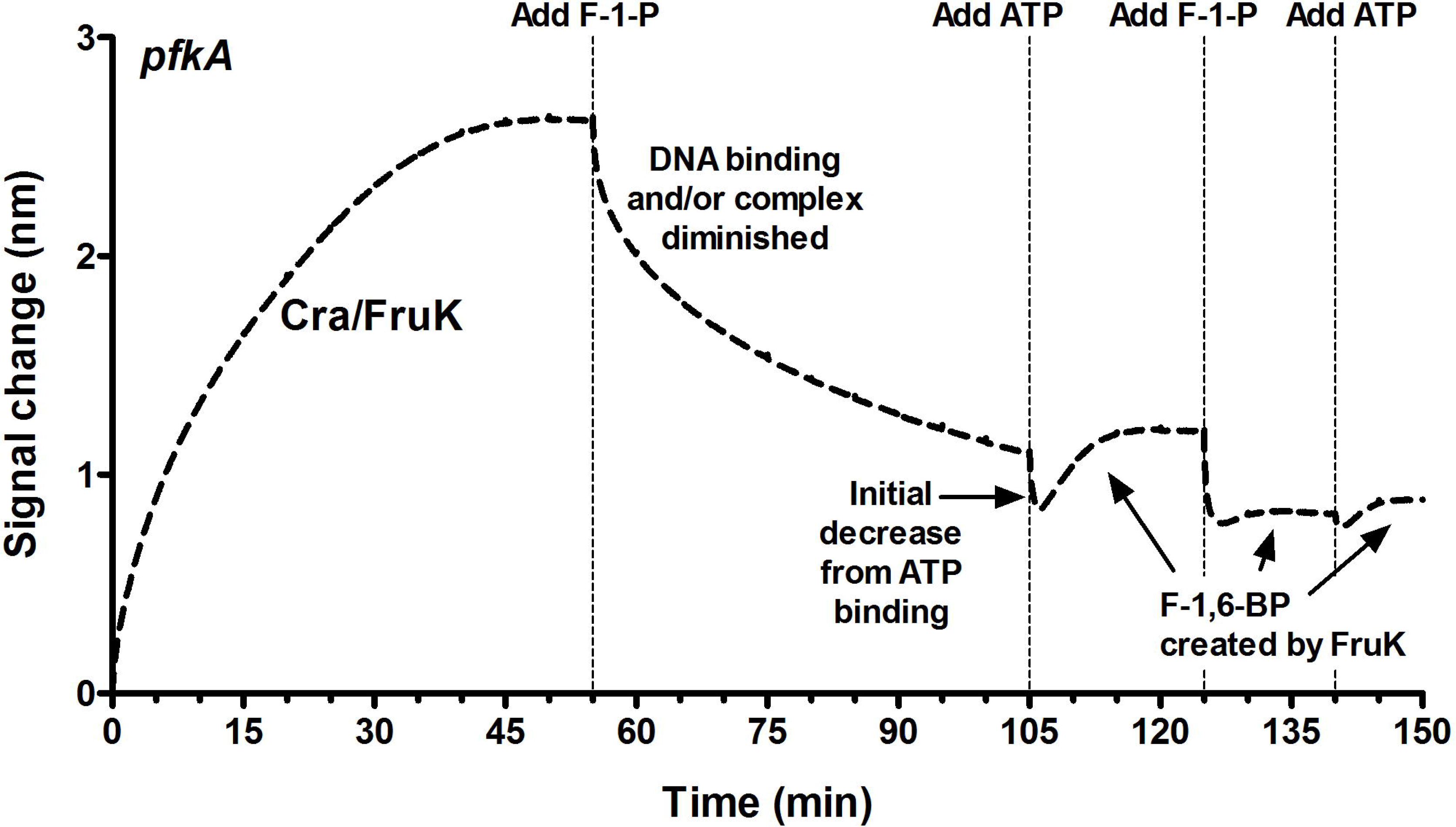
Phosphorylation of F-1-P monitored with Cra/FruK binding to DNA. Biotinylated *pfkA* DNA was immobilized to streptavidin tips from an 80 nM solution. Cra (50 nM) was pre-equilibrated with 200 nM FruK and bound to the DNA. Addition of the inducer F-1-P (1 mM) diminished the signal, as did the initial addition of 3 mM ATP. However, as FruK catalysis generated F-1,6-BP from these substrates, the signal change reversed and began to increase. Similar effects were observed for a second cycle of F-1-P and ATP additions, with catalysis apparently occurring after each step and reversing the signal change. We attribute the diminished response of these later steps to the build-up of F-1-P (either uncatalyzed or *via* the reverse reaction), which binds Cra (and perhaps Cra-FruK) with µM affinity and diminishes DNA binding.

To detect the reverse enzymatic reaction, we added F-1,6-BP and then ADP to DNA-bound Cra/FruK. The second addition caused a large decrease in the biolayer interferometry signal (Figure 11), with a final amplitude very similar to that of [Cra + F-1-P]. This result was consistent with enzymatic conversion of F-1,6-BP to F-1-P, followed by F-1-P binding to and induction of Cra to diminish DNA binding. The effect occurred for even the tightest *fruB* operator (Supplementary Figure 7) at the decreased concentration of 57 nM FruK and the physiological concentration of 0.3 mM ADP (Figure 11B). As a control, we performed similar experiments with 1 mM cyclic AMP with F-1,6-BP; no effect was observed (data not shown).

**Figure 11.**
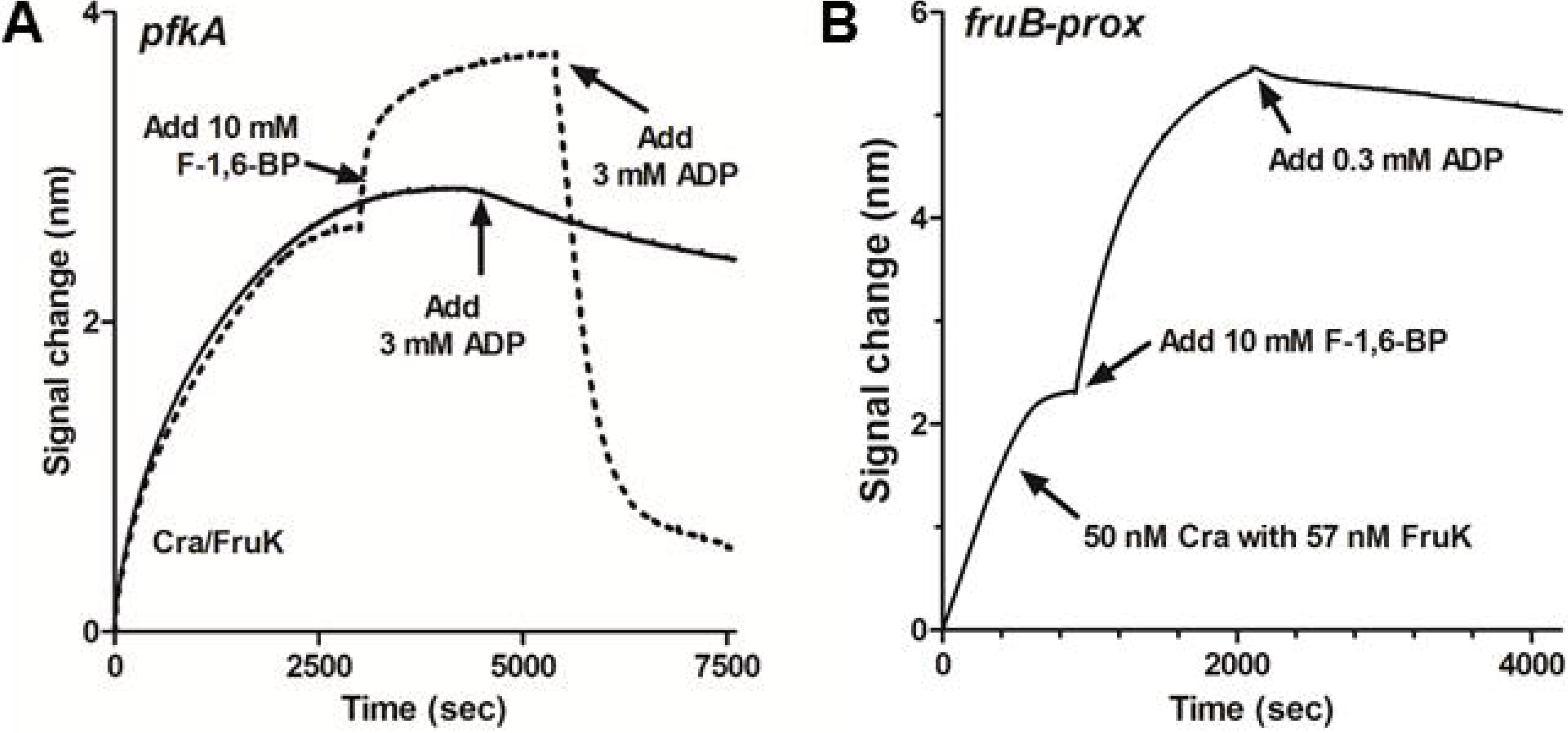
De-phosphorylation of F-1,6-BP monitored with Cra/FruK binding to DNA. In both experiments shown, 50 nM Cra was pre-equilibrated with 200 nM FruK and bound to the DNA indicated (80 nM solution). (A) Addition of the F-1,6-BP (10 mM) enhanced the signal, as expected (dotted line). When 3 mM ADP was added after F-1,6-BP, the signal dramatically decreased (dotted line), which we attribute to the catalytic creation of the strong inducer F-1-P. In the control experiment, addition of 3 mM ADP slightly diminished the signal (solid line). (B) Addition of 10 mM F-1,6-BP enhanced formation of the Cra/FruK:DNA complex even at the low concentration of 57 nM FruK. Addition of 0.3 mM ADP showed a slight decrease. Control experiments with Cra/FruK:DNA with 0.3 mM ADP added showed no change (not shown).

Next, we directly monitored the effects of varied substrate concentration and order of addition on the protein-DNA complex. To that end, we pre-equilibrated Cra/FruK with immobilized DNA then subsequently added small molecules. Experiments used the *fruB* operator DNA, 200 nM FruK, and the physiological concentration of ADP (0.3 mM). As was true for 10 mM F-1,6-BP, when 1 mM F-1,6-BP was the first substrate added, the signal increased and the addition of the 0.3 mM ADP initiated the decreasing signal. When 0.3 mM ADP was the first substrate added to the Cra/FruK:DNA complex, the signal did not change appreciably until F-1,6-BP was added. In the presence of 0.3 mM ADP, even 0.013 mM F-1,6-BP was sufficient to initiate the decreasing signal.

We were surprised that no concentration dependence was readily observed over the range of 0.013 mM to 20 mM F-1,6-BP. (The different decay rates seen in panels A and B of Figure 11 are best explained by the different FruK concentrations.) At least two scenarios can explain this behavior: (i) The K_m_ of FruK for F-1,6-BP is below 0.013 mM; note that 0.013 mM is still above the micromolar concentration of F-1-P needed to induce Cra. Or (ii) FruK catalysis of the reverse reaction is much faster than dissociation of the Cra/FruK:*fruB* complex, which means that the latter is a rate limiting step and cannot be used to quantify the enzymatic assay. In addition, this assay does not distinguish whether the free or Cra/DNA-bound FruK (or both) is the catalytically active species.

## Discussion

*E. coli* Cra is routinely referred to as a “master regulator” for switching between glycolytic and gluconeogenic metabolism. However, as noted in the introduction, the mechanism of such a metabolic switch has been difficult to fully rationalize using the previously known parameters of (i) strong inducer F-1-P only present when fructose is an available nutrient, and (ii) weak inducer F-1,6-BP derived from the metabolism of many substrates. The results reported here provide several new features that make the model of Cra regulation more complete. Specifically, this work provides evidence that: (i) Cra forms a complex with the FruK protein that may have enhanced DNA binding properties. (ii) Deleting the gene for FruK impairs the switch from rich media to glycerol utilization, suggesting a broad role for this protein in metabolic regulation. (iii) Some aspect of Cra/FruK:DNA complex formation is enhanced by the presence of F-1,6-BP. (iv) The reverse FruK reaction can occur to generate F-1-P from F-1,6-BP.

To formulate a biological model for the Cra/FruK complex, we have integrated these observations with two here-to-fore unexplained observations by Molchanova et al. (Molchanova *et al.*, 1992). First, these authors showed that *fruK* mutant strains of *E. coli* are unable to grow on lactate^a^, which they ascribed to a decrease in PEP-synthase (Molchanova *et al.*, 1992). Second, they showed that a strain with double *fruK*, *cra* knock-outs caused even further decreases in PEP synthase. Notably, lactate is gluconeogenic (Hua *et al.*, 2007), and many genes in the gluconeogenesis pathway (including PEP synthase) are activated by Cra (Chin *et al.*, 1989). We propose that Molchanova’s results are explained by FruK and/or FruK/F-1-6-BP enhancement of Cra activation. (F-1,6-BP levels are not yet known for lactate carbon source.) This is consistent with our results showing that Cra/FruK and Cra/FruK/F-1,6-BP may better bind DNA than does Cra alone.

Loss of enhanced Cra-FruK activation can also explain the increased glycerol lag time we observed for the Δ*fruK* strain: Glycerol can either feed into the TCA cycle or be gluconeogenic (Torres *et al.*, 1997) and both pathways are activated by Cra (Saier & Ramseier, 1996, Ramseier *et al.*, 1995). When the carbon source switched to glycerol, the wild-type strain would have full Cra/FruK/F-1,6-BP activation of both pathways. (Note in Table 1 that glycerol growth conditions produce ∼5.9 mM F-1,6-BP; biochemical experiments showed enhanced Cra/FruK:DNA binding over the range of 1 to 10 mM F-1,6-BP.) In contrast, the Δ*fruK* strain would have only the lower Cra-only activation, which would in turn lead to more modest enzyme expression. (Even if the Δ*fruK* grown on glycerol generates the same level of F-1,6-BP as wild-type strains, the direct effect of 5 mM F-1,6-BP does not diminish Cra-DNA as well as F-1-P (Ramseier *et al.*, 1993).) However, enough enzyme could eventually accumulate in the Δ*fruK* strain to allow the bacteria to resume logarithmic growth.

The potential biological implications of the reverse FruK reaction are more complicated to resolve. Loss of this reaction does *not* appear to contribute to the lactate or glycerol growth defects of Δ*fruK* strains: In wild-type *E. coli*, the reverse reaction would work against formation of the Cra/FruK:DNA complex (generating F-1P that would out-compete F-1,6-BP) and thereby diminish DNA binding. In Δ*fruK*, loss of the reverse reaction should enhance DNA binding and activation of gluconeogenic genes. These predictions are opposite to the results described above.

Nevertheless, the reverse reaction occurs at concentrations within or well below the physiological range for *E. coli* (Table 1) (Bennett *et al.*, 2009). Since our experiments were carried out with excess FruK, they do not resolve whether the Cra-free or bound form of FruK catalyzes the reverse reaction. One possibility is that reverse reactions of the Cra-free FruK play a role somewhere else in the cell. Another possibility is that the reverse FruK reaction acts as a missing component of the proposed (Kochanowski *et al.*, 2013, Usuda *et al.*, 2010) flux sensor: The F-1,6-BP that arises from the metabolism of many sugars could be converted to the strong inducer F-1-P.

Another possible source of the flux sensor is the levels of FruK protein itself. The gene for FruK is repressed by Cra: As nutrient conditions change the Cra induction status, FruK levels must cycle between high and low concentrations. In turn, the amount of Cra in complex with FruK would change, allowing any concomitant changes in DNA binding. A range of possible *in vivo* FruK concentrations can be deduced from our experiments: The FruK concentration is never zero (even when Cra is uninduced), otherwise endogenously encoded FruK would not co-purify with over-expressed Cra. Conversely, when Cra was induced for pulldown assays (Figure 2), endogenously encoded FruK was expressed at about the same level as the repressor protein, which we previously determined is >2500 copies per cell (Tungtur *et al.*, 2011).

Several other parameters are still needed to fully model the effects of the Cra/FruK complex on the regulation of *E.coli* metabolism. One question is the stoichiometry of the Cra/FruK complex. An intriguing possibility is that FruK binding promotes Cra tetramerization. In our hands, Cra eluted as a dimer from the phosphocellulose purification column, as benchmarked against the tetrameric, dimeric (Swint-Kruse *et al.*, 2005), and monomeric (Chen & Matthews, 1992) forms of homolog LacI. In contrast, Cozzone and colleagues previously reported tetramerization of His-tagged Cra (Cortay *et al.*, 1994). However, in the earlier experiment, the presence of co-purifying FruK would not have been apparent. (We have only been able to resolve the two proteins on one type of gel; see Experimental Procedures). These different outcomes may very well indicate that Cra quaternary structure is modulated by interaction with FruK.

A second set of missing parameters pertain to the effects of Cra/FruK on varied DNA sequences. The Cra regulon comprises 24 operons regulated by individual operator sequences (Ravcheev *et al.*, 2014b), and Cra/FruK complex formation could provide a mechanism for their differential regulation. Notably, the current work did *not* show a difference in complex formation when binding single DNA operators from three functionally-distinct operons (*fruBAK*, *aceB*, and *pfkA*). However, the *fruBAK* operon has two strong operator binding sites. If the FruK interaction were to alter Cra quaternary structure, tetrameric Cra could loop two binding sites of an operon and enhance repression of the relevant genes, in parallel to tetrameric LacI (Swint-Kruse & Matthews, 2009). In support of this idea, we previously documented that the LLhF chimera shows repression consistent with *in vivo* looping of the *lac* operon (Meinhardt *et al.*, 2012).

Finally, we note the intriguing parallel between the Cra/FruK/F-1,6-BP response and the LacI homolog CcpA. CcpA is a central metabolic regulator of Gram positive bacteria that is regulated by both protein-protein interactions (with HPr and CPr) and by F-1,6-BP and glucose-6-phosphate binding to the regulatory domain (Richardson *et al.*, 2015, Seidel *et al.*, 2005, Schumacher *et al.*, 2007). The parallels between Cra and CcpA regulation point to the universal, central role played by the F-1,6-BP metabolite. A number of other examples demonstrate the importance of central metabolism to pathogenicity. For example, sugar use must be balanced between energy generation and biosynthesis of cellular components, such as cell wall/biofilm construction (Forsberg *et al.*, 2012, Kawada-Matsuo *et al.*, 2012, Zhang *et al.*, 2012, Willenborg *et al.*, 2011). In a second example, enteric pathogens must sense variations in nutrient availability as they move through the gastrointestinal tract and identify the proper region for infection (Gore & Payne, 2010, Njoroge *et al.*, 2012). The Cra/FruK interaction in y-proteobacteria such as *E.coli* may provide a unique way to target this biology without disturbing metabolism of the mammalian host or beneficial microbes. With the impending doom of antibiotic resistance, this may allow us to re-gain the evolutionary upper hand.

## Experimental Procedures

### Plasmid construction

Creation of the pHG165a plasmid encoding the LacI:Cra chimera LLhF was reported in 2012 (Meinhardt *et al.*, 2012). The low copy pHG165 plasmid (Stewart *et al.*, 1986) uses the *lacl*^q^ promoter for constitutive gene expression and carries an ampicillin resistance gene. To carry out experiments with wild-type Cra, the gene was amplified from Hfr(PO1), *lacI22, λ-, e14-, relA1, spoT1, thiE1 E. coli* (strain name 3.300; *E.coli* Genetic Stock Center, Yale University) using the primers 5'-GGTCTCCAGGTGTGAAACTGGATGAAATCGC-3' and 5'-GAGCTCTTAGCTACGGCTGAGCACGCCGCGG-3'. The gene was then subcloned into the pHG165a plasmid using the procedure of Jones (Jones, 1994). A plasmid carrying the *fruK* gene (pET-3a) was a gift from the Van Schaftingen laboratory at the Université Catholique de Louvain, Belgium (Veiga-da-Cunha *et al.*, 2000).

### Growth curves of WT and AfruK

To detect the biological outcomes of *fruK* deletion, we performed growth assays with the parent F-, *Δ(araD-araB)567, ΔlacZ4787*(::rrnB-3)*, λ-, rph-1, Δ(rhaD-rhaB)568, hsdR514* (strain name BW25113) and knockout F-, *Δ(araD-araB)567, ΔlacZ4787*(::rrnB-3)*, λ-, ΔfruK724::kan, rph-1, Δ(rhaD-rhaB)568, hsdR514* (strain name JW2155-1) *E. coli* strains from the Keio collection (Baba *et al.*, 2006); both were obtained from the *E. coli* Genetic Stock Center at Yale University (New Haven, CT). In order to confirm the disruption of *fruK* in JW2155-1, the following primers were designed to flank the *fruK* gene (NCBI gene ID: BW25113_RS11315; WP_000091263.1): fwd 5’-AGCAGCAGCGTTTTCATTATG-3’; rev 5’-GACGCTATCGCTGCTGG-3’. NEB Q5® High-Fidelity polymerase (Ipswich, MA) was used in a standard PCR reaction and the transposon disruption of *fruK* resulting amplicons was confirmed by sequencing (ACGT, Germantown, MD).

The growth assay protocol was modified from that of Hall *et al.* (Hall *et al.*, 2014). Briefly, both the wild-type and *ΔfruK* strains were grown in 10 mL rich media (Luria broth) until late-log phase (OD600∼1.5 in a cuvette with 1 cm path length). Cultures were then centrifuged at 1800хg for 15 mins, decanted, and resuspended in 1mL “M9” media (182 mM sodium phosphate, 110 mM potassium phosphate monobasic, 93 mM ammonium chloride, 42 mM sodium chloride, 1 mM magnesium sulfate, 0.00005% thiamine, pH 7.4) containing 9 mM MgCl_2_ and 0.2% w/v of glucose, glycerol, or fructose as the carbon source, to a starting OD_600_ of ∼0.1. The additional MgCl_2_ was to facilitate FruK activity and complex formation, which both require Mg^2+^. Subsequently, 100µL of resuspended bacteria were aliquoted into quadruplicate wells of a 96-well plate (Becton Dickinson, NJ).

Bacterial growth was then monitored using either a BioTek Synergy2 microplate reader (Winooski, VT) or a Tecan M200 pro (Switzerland) with data collection via either Gen5 or Magellan data analysis software. During these experiments, the plate was kept at 37 °C with shaking (198.4 rpm); the OD_600_ was measured every 15 mins for 24 hours. Media-only wells served as blanks and were subtracted from bacteria-containing wells. Doubling times were calculated for the logarithmic growth phase using the inverse slope of log_2_ (OD_600_) values. Data shown are the mean and standard deviation of quadruplicates samples from each of four independent experiments; unpaired t-tests using were used to assess differences in growth rates. The growth delay time as the WT and *ΔfruK* strains transitioned from rich to minimal media (t_delay_) was calculated as the mean time required for bacteria to reach mid-log phase (OD_600_ 0.15). Data shown are the mean and standard deviation of quadruplicates samples from each of four independent experiments; paired t-tests were applied to determine statistical differences in growth delay. All calculations were performed with GraphPad Prism v7.0.

### Biotinylated DNA

For pulldown assays and biolayer interferometry, full-length biotinylated DNA was either purchased from Integrated DNA Technologies (Skokie, Illinois) or amplified in a thermocylcer using a short oligo of DNA containing a biotin tag and primers encoding the desired DNA operator sequences. For the latter, the single stranded DNA oligos were mixed in a 1:1 ratio in Taq polymerase buffer that contained 2.5 mM dNTPs and 2.5 units Taq Polymerase (New England Biolabs). The mixture was heated to 95° C for 1 minute, then the temperature was reduced 5° C per minute until the temperature reached 35° C. Product formation was verified via electrophoresis on a 2.5% agarose gel.

### Sources for F-1-P, F-1,6-BP, ATP, and ADP and ion exchange of F-1-P

Early enzyme assays and DNA binding experiments used the highly soluble sodium salt of F-1-P obtained from Sigma-Aldrich (St. Louis, MO). When that compound was discontinued, we purchased the less-soluble barium salt of F-1-P from the same company. To obtain the F-1-P concentrations necessary for enzymatic assays, we performed ion exchange chromatography of the barium salt using a Dowex 50WX4 (Sigma) column pre-charged with 2 N HCl and then equilibrated in ddH_2_O. The barium F-1-P was passed through this column several times and the pH of the final eluant was adjusted to 7 with NaOH. Eluted F-1-P was lyophilized and used to make saturated stock solutions in ddH_2_O. Concentrations of the stock solutions were determined by stoichiometric implementations of the enzyme assay (see below).

The trisodium salt of D-fructose-1,6-bisphosphate (F-1,6-BP) was purchased from Sigma-Aldrich (St. Louis, MO). ATP and ADP were purchased from Sigma-Aldrich (St. Louis, MO). For experiments, these nucleotides were dissolved in water and the pH was adjusted to 7.0.

### DNA pull-down assays

For pull-down assays, plasmids encoding either the LLhF chimera or wild-type Cra were transformed into the *E.coli* strain 3.300. Cultures were grown at 37°C to saturation in 3 mL of MOPS minimal media (Teknova, Hollister, CA: 40 mM morpholinopropanesulfonic acid, 10 mM NH_4_Cl, 4 mM tricine, 50 mM NaCl, and other trace metals listed for product number M2101) supplemented with 0.8% glycerol, 1.32 mM dibasic potassium phosphate, 10 mM NaHCO_3_, 0.2% casamino acids, 0.0025% thiamine and 100 µg/ml ampicillin (Neidhardt *et al.*, 1974, Bhende & Egan, 1999). Cells were pelleted by centrifugation, resuspended in 100 µL breaking buffer (0.2 M Tris-HCl, 0.2 M KCl, 0.01 M MgCl_2_, 1 mM dithiothreitol [DTT], 5% glucose, pH 7.6) and lysed via freeze-thaw with 40 µL of 5 mg/ml lysozyme. Crude cell lysates were clarified by centrifugation. Ten (10) µL supernatant were incubated with biotinylated DNA that was immobilized to streptavidin magnetic beads (1 pmol DNA to 1 µL beads; New England Biolabs, Ipswich, MA), as described in Meinhardt *et al.* (Meinhardt & Swint-Kruse, 2008). Sequences of the biotinylated DNA operators (Integrated DNA technologies, Coralville, IA) used here and in biolayer interferometry (below) are listed in Table 2. After washing in FB buffer (10 mM Tris HCl, 150 mM KCl, 10 mM EDTA, 5% DMSO, 0.3 mM DTT, pH 7.4), the protein-DNA-beads were resuspended in 15 µL of denaturing buffer (10 µL 20% sodium dodecyl sulfate plus 5 µL 1M dithiothreitol [DTT]). Adherent proteins were visualized with SDS-PAGE.

**Table 2.**
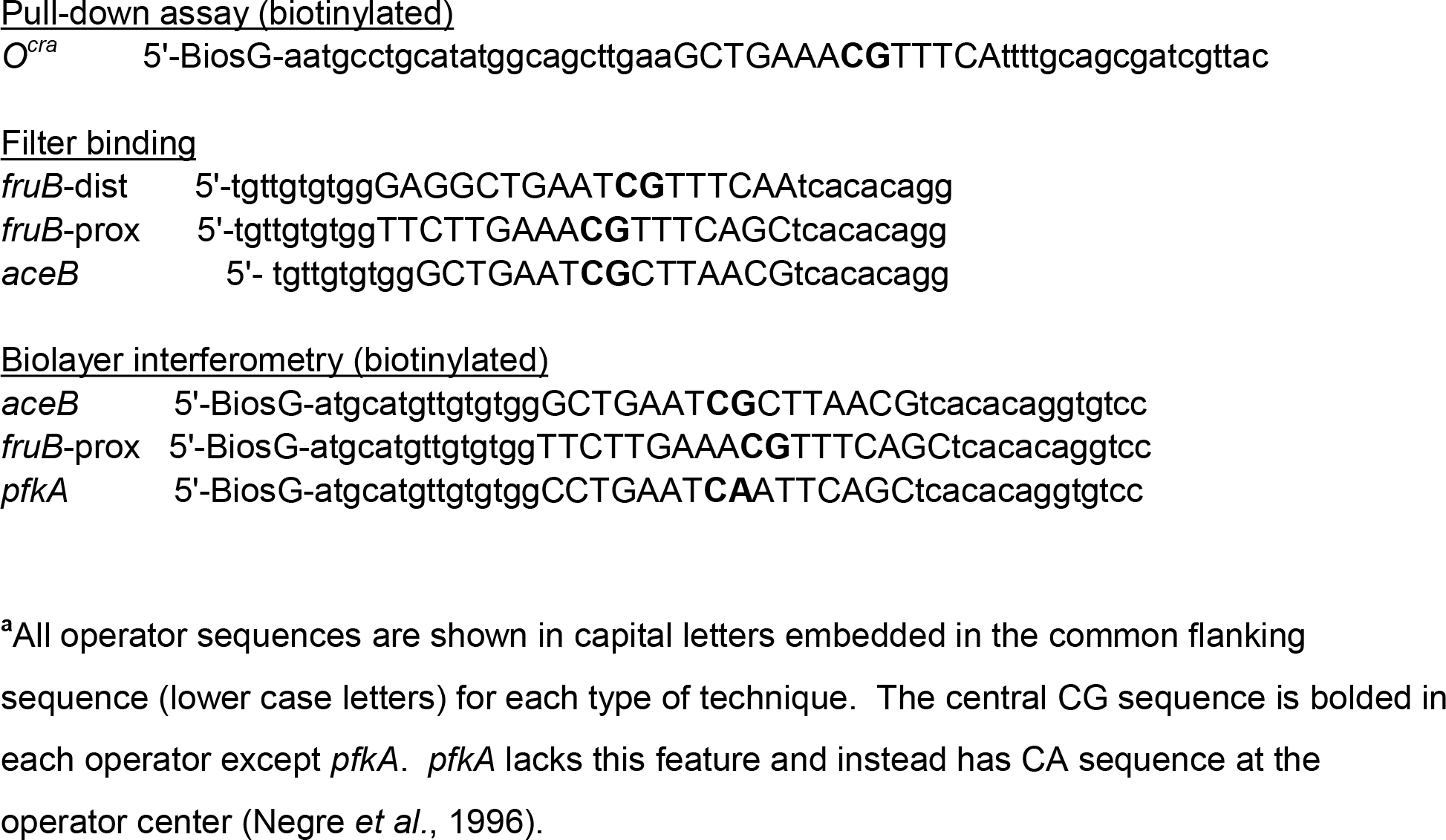
Operator sequences used in Cra/FruK experiments.^a^

In pull-down assays with either LLhF or Cra, a second, slightly smaller protein band was visible. To date, the only gel that resolves the repressor from the second band is the Phast-gel 10-15% gradient, (GE Healthcare, Piscataway, NJ; catalogue #: 17-0540-01).

### Mass Spectrometry

For protein identification, the two bands from the pull-down gel were excised and sent to the Mass Spectrometry Core Laboratory at KUMC for analysis; this confirmed the identity of LLhF in the top band and identified the second band as FruK. To that end, the cut-out bands were subjected to trypsin digestion and submitted for mass spectrometry analyses. Results were analyzed using the Sequest algorithm to search an *E. coli* protein database derived from the NCBInr repository on February 2, 2011. Trypsin specificity was requested, with a maximum of two missed cleavages. Oxidation of methionine, deamidation of asparagine and glutamine, and carboxymethylation of cysteine (defined as a fixed modification) were defined as variable modifications. Statistical criteria for peptide identification included: a minimum XC of 1.8, 2.5 or 3.5 for a peptide ion with a charge of 1, 2, or 3, respectively, a minimum Delta Cn of 0.01, and a probability of 0.001. For protein identification, a minimum of two unique peptides was required. Results are shown in Supplementary Table 1.

### Cra and LLhF purification

Cra: The plasmid encoding wild-type Cra was constitutively expressed in *E.coli* B *omp*T, 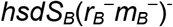, *gal, dcm, lac* cells (strain name BLIM (Wycuff & Matthews, 2000)). All cultures were grown in the presence of 50 mg/mL ampicillin. Cells were grown overnight at 37°C in 3 mL 2xYT media, of which 500 µL was subsequently used to inoculate 50 mL 2xYT media. The 50 mL media was grown for 8 hours at 37°C then used to inoculate 6 L of 2xYTmedia (1 L media per 2 L Erlenmeyer flask). The large cultures were grown 19-20 hours at 37°C with shaking at 220 rpm.

The protocol to purify Cra is similar to that used to purify lactose repressor protein (LacI) (Falcon & Matthews, 2001, Swint-Kruse *et al.*, 2003, Swint-Kruse *et al.*, 2005, Taraban *et al.*, 2008, Zhan *et al.*, 2006). After pelleting by centrifugation, the cells grown in 6 L 2xYT media were resuspended in ∼50 mL breaking buffer (0.2 M Tris-HCl, 0.2 M KCl, 0.01 M MgCl_2_, 1 mM DTT, 5% glucose, pH 7.6) with 20 mg lysozyme and half of a crushed Pierce Protease Inhibitor Tab (Thermo Scientific, Rockford, IL) and frozen at −20°C. All subsequent purification steps were carried out on ice or at 4°C. Cell lysis was accomplished by thawing the cell pellet on ice with breaking buffer added to a final volume of 150 mL and another crushed protease tab. At this step, the addition of 120 mg extra lysozyme was critical for chaperoning the Cra through ammonium sulfate precipitation, which was an unexpected activity of lysozyme. After lysis, 240 µL of 80 mg/ml DNase I from bovine pancreas (Roche Diagnostics, Mannheim, Germany) was added along with MgCl_2_ to a final concentration of 30 mM. The crude cell extract was cleared by centrifugation at 8,000 rpm for 50 min at 4°C prior to 23% ammonium sulfate precipitation (Fisher Scientific, Fair Lawn, NJ).

The technical aspects of ammonium sulfate precipitation were very important. The chemical must be finely ground and lightly salted over the whole surface of the solution (generally 180-200 mL in a 400 mL beaker) at a rate so slow that crystals never reach the bottom of the flask. The mixture was allowed to precipitate for 30 minutes at 4°C with slow stirring prior to centrifugation at 8000rpm for 40 min and 4°C.

After precipitation, the pellet was gently resuspended in 50 mL cold 0.09 M KP buffer (KP buffer comprises potassium phosphate at the indicated molarity, 5% glucose, 0.3 mM DTT, pH 7.5) and dialyzed in regenerated cellulose Fisherbrand Dialysis Tubing (MWCO 12,000-14,000) against 1 L of cold 0.09 M KP buffer for 30 minutes, with three buffer changes. The dialysate was cleared by centrifugation, diluted 1:6 with 0.09 M KP buffer, and loaded onto a phosphocellulose column (Whatman P11) that was pre-equilibrated in 0.09 M KP buffer. The Cra protein eluted shortly after the beginning of a 0.09 M to 0.2 M KP buffer gradient, which was benchmarked against LacI variants as behaving like a dimer (Swint-Kruse *et al.*, 2005, Chen & Matthews, 1992). The pre-column dilution (1:6 in 0.09 KP buffer) was key to good purity; without it, Cra eluted from the phosphocellulose column much later and was contaminated with lysozyme and other proteins. Purified Cra (Supplementary Figure 2) was aliquoted and stored at −80°C. Protein concentration was determined by Bradford assays (Bio-Rad Laboratories, Inc. Hercules, CA) and was generally 0.1-0.2 mg/ml (4.2 µM, assuming 100% Cra).

LLhF: The LacI:Cra chimera LLhF was purified similarly to wild-type Cra. However, when expressed from the low-copy pHG165a plasmid, the chimera purified as a complex with FruK in a nearly 1:1 ratio. The fact that the LLhF-FruK complex persisted through all steps of the purification protocol indicates extremely tight binding between the two proteins. Enzyme assays were used to confirm the identity of FruK as the contaminating band. We cannot currently discriminate (i) whether LLhF binds more tightly to FruK than does Cra, or (ii) whether more FruK is expressed in these cells because LLhF is dominant negative heterodimer over wild-type Cra. The amount of LLhF/FruK complex was significantly diminished (but not abolished) by cloning and constitutively expressing the chimera on the high copy pLS1 plasmid (Swint-Kruse *et al.*, 2001)(Supplementary Figure 2), which also uses the *lacl*^q^ promoter for constitutive gene expression and carries an ampicillin resistance gene. LLhF was further purified by the following steps: To lower the ionic strength of the buffer, LLhF protein was dialyzed against a buffer containing 5 mM HEPES, 5 mM KCl, 0.1 mM EDTA, 10 mM MgCl2, 0.3 mM DTT, pH 7; concentrated in Vivaspin MWCO 10 kDa at 4 °C; and loaded onto a GE HiPrep 16/10 Heparin column equilibrated in the same buffer. LLhF was eluted with a gradient of the loading buffer and a high salt buffer comprising 50 mM HEPES, 500 mM KCl, 0.1 mM EDTA, 10 mM MgCl2, 0.3 mM DTT, pH 7. The heparin-purified LLhF was used for biolayer interferometry experiments.

### FruK purification

For protein expression, the FruK-pET-3a plasmid (Veiga-da-Cunha *et al.*, 2000) was transformed into *E.coli* B F^−^ *ompTgal dcm lon* 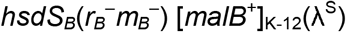 cells (strain name BL21). Freshly-made 3 mL aliquots of 2xTY media were inoculated and grown overnight at 37°C. Six (6) mL of these cultures were subsequently used to inoculate each of three 2 L flasks containing 1 L M9 media (211 mM Na_2_HPO_4_, 110 mM KH_2_PO_4_, 93.4 mM NH_4_Cl, 42.8 mM NaCl,135 µM CaCl_2_, 1 mM MgSO_4_ * 7H_2_O, 0.2% Glucose, 50 ug/ml ampicillin, 0.00005% vitamin B1, 0.1% casamino acids, pH 7.4). Based on the previous publication (Veiga-da-Cunha *et al.*, 2000), we also attempted protein expression in rich media, but minimal media gave superior FruK concentration and purity as assessed by SDS-PAGE.

We also obtained better expression/purification by inducing FruK expression at the end of the previously published induction window of 0.4-0.6 OD_600_: We grew 1 L M9 media for 3-5 hours at 37°C until OD_600_ reached 0.6, then induced with 1 mM isopropyl-1-thio-β-D-galactopyranoside (IPTG). Induced cultures were incubated at 22°C with shaking at 220 rpm for an additional 44-45 hours. Cells were pelleted at 7,000 rpm for 15 min and resuspended in 50 mL of FruK breaking buffer (20 mM potassium phosphate, 5mM EDTA, 1 mg/mL lysozyme, 1 Pierce Protease Inhibitor Tablet (Thermo Scientific, Rockford, IL], 1mM DTT, pH 7.4). The crude cell lysate was cleared by centrifugation, and protein purification proceeded as described (Veiga-da-Cunha *et al.*, 2000) using a Sephadex G25 column for the desalting step and DEAE-sepharose for the anion exchange.

After the final column, the FruK fractions were brought to 20% glycerol prior to freezing at −80°C. Protein concentrations were generally around 0.065 mg/ml, as determined by Bio-Rad assays. We attempted to remove the glycerol by dialysis prior to enzyme assays, but found that without this osmolyte, the FruK enzymatic activity decayed over the course of repeated assays (30 min - 1 hour). A broad survey of other freezing methods, temperatures, and using glucose as a cryoprotectant showed that glycerol was superior for maintaining enzymatic activity. Thus, the enzyme and DNA binding assays used FruK with 20% glycerol.

### FruK enzyme assay

The published assay for FruK activity (Veiga-da-Cunha *et al.*, 2000) was successfully repeated, but we found that the commercial aldolase used as a coupling enzyme had inconsistent and sometimes quite high activity towards F-1-P. Therefore, we devised a new assay by coupling the FruK reaction to that of pyruvate kinase from rabbit muscle (Roche, Mannheim, Germany) and lactate dehydrogenase (Calzyme Laboratories, San Luis Obispo, CA). Assays used 5.23 U/ml lactate dehydrogenase and 10 mg/ml pyruvate kinase, following the example of pyruvate kinase studies (Fenton & Alontaga, 2009). Other reagents included phosphoenol pyruvate (PEP; 10 mM; Chem-Impex, International, Wood Dale, IL), ATP (Chem-Impex; 3mM, unless otherwise indicated), NADH (Sigma Chemical Co; 0.18 mM), and F-1-P. The reaction rate for FruK was monitored by the depletion of NADH at 340 nm in a SPECTRAmax PLUS 384 at 30°C using a UV-Star flat bottom 96-well plate (Greiner Bio-One, Monroe, NC). The reactions were initiated by the addition of ATP, and initial velocities were determined as a function of F-1-P concentration and used to estimate values of K_m_ with the program GraphPad Prism 5 (GraphPad Software Inc., La Jolla, CA).

Two different assay buffers were compared: (i) that of Viega de Cunha and colleagues (Veiga-da-Cunha *et al.*, 2000) (20 mM Hepes, pH 7.1, 5 mM MgCl_2_, and 50 mM KCl, with 2.5 mM ATP and 0.15 mM NADH) and (ii) that used for pyruvate kinase assays (Fenton & Alontaga, 2009) (“PYK buffer”: 50 mM Hepes, 7.0 pH, 10 mM MgCl_2_, and 0.1 mM KCl, with 3 mM ATP and 0.335 mM NADH). Of the two, the PYK buffer is more similar to DNA binding buffer used in filter binding assays (“FBB”, see below). We observed no noticeable difference in enzyme activity between the two buffers and therefore continued with the PYK buffer.

A stoichiometric implementation of the PYK-coupled assay was used to determine the concentration of the ion-exchanged F-1-P. This assay used a high concentration of FruK so that catalysis of the F-1-P substrate was quickly completed. The change in amplitude at 340 nm was used to determine the total amount of NADH consumed during the reaction; this value corresponds 1:1 with the F-1-P concentration.

### Cra-DNA binding assays

Cra-DNA binding assays were carried out in the absence and presence of added, purified FruK. For the FruK-free experiments, we were concerned about the contaminating FruK that co-purified with both Cra and LLhF. However, the repressor proteins were diluted ≥10-fold, to a final concentration of 100 nM or less for DNA binding assays. This appeared to dilute contaminating FruK to levels below those required for significant complex formation, as evidenced by the effects observed when high concentrations of FruK were added to the Cra-DNA binding reactions. We did find that some Cra preparations with low concentrations after purification (from the edges of the elution peak) had a higher percent FruK contamination: When these protein samples were concentrated for biolayer interferometry assays, F-1,6-BP additions on “Cra” had the outcome expected for the Cra/FruK complex.

One type of DNA-binding assay used the strategy of nitrocellulose filter binding (Amersham Protran 0.45 µm, GE Healthcare Life Sciences) with ^32^P-labeled operator DNA (Table 2), as described previously (Swint-Kruse *et al.*, 2003, Zhan *et al.*, 2006, Falcon & Matthews, 2001, Riggs *et al.*, 1968, Wong & Lohman, 1993). Experiments used two different buffers: (i) “FBB” of the LacI DNA binding assays (10 mM Tris, 150mM KCl, 10 mM EDTA, 0.3 mM DTT, pH 7.4, 0.1 mg/ml bovine serum albumin [BSA]), and (ii) the enzyme assay buffer of Viega de Cunha (listed above). The different buffers had little or no impact on K_d_ values, which was especially surprising given the variation in salt concentrations. Control experiments with varied salt and pH indicated that the differences in these two parameters directly offset each other, so that the two buffers appeared equivalent. For stoichiometric assays, the DNA concentration was fixed at least 10-fold greater than the value of K_d_; for equilibrium assays, the DNA concentration was fixed at least 10-fold less than the value of K_d_ (Swint-Kruse & Matthews, 2004). Experiments were repeated multiple times, using at least two different preparations of protein and both buffers.

A second type of DNA binding assay monitored changes in the interferometry of biosensor tips (Concepcion *et al.*, 2009) (BLItz; Forté Bio, Pall Life Sciences, Menlo Park, CA) in the presence of Cra, FruK, and various small molecules. In the current experiments, probe tips were derivatized with streptavidin to which various biotinylated DNA sequences were attached (Table 2). To that end, tips were first hydrated for at least 10 minutes in the DNA immobilization buffer, loaded with biotinylated DNA (120 sec), washed in immobilization buffer (30 sec), and then dipped into pre-equilibrated solutions of protein(s) and any small molecules. Immobilization buffer (50 mM Hepes, 0.1 M KCl, 0.1 mM EDTA, 10 mM MgCl_2_, pH 7.0, 0.3 mM DTT) was similar to the PYK buffer used in the FruK enzyme assay.

Most experiments used Streptavidin-derivatized tips (Forté Bio, Pall Life Sciences, Menlo Park, CA), which had low nonspecific binding to either Cra or FruK in the absence of DNA; however tip-to-tip amplitudes were less reproducible. For later experiments, a second “High-Precision Streptavidin” tip became available from Forté-Bio that had better tip-to-tip reproducibility but greater nonspecific binding of Cra and FruK in the absence of DNA. The latter was blocked by adding 0.1 mg/mL BSA to the initial hydration buffer.

For DNA loading, immobilization buffer contained 20-240 nM biotinylated DNA; most experiments used 80 nM. Protein solutions were kept on ice until added to final reaction tube, which was then allowed to warm to room temperature (∼15 min). (Biolayer interferometry signals are highly temperature dependent.) All biolayer interferometry steps were carried out with shaking at 2,200 rpm. Resulting signal changes were monitored as a function of time. In other experiments, the Cra or Cra/FruK proteins were first equilibrated with the immobilized DNA and either buffer or solutions of the relevant small molecules were added at later times.

We first benchmarked the biolayer interferometry assays using the well-characterized DNA-binding events of LacI (*e.g.* (Swint-Kruse *et al.*, 2003, Zhan *et al.*, 2006, Falcon & Matthews, 2001)) and the LacI:PurR chimera “LLhP” (Zhan *et al.*, 2008) (Supplementary Figures 3-6; additional references cited in the Supplement are listed here (Oehler *et al.*, 1990, Pfahl *et al.*, 1979, Gilbert & Maxam, 1973)); these proteins are homologs of Cra and LLhF (Swint-Kruse & Matthews, 2009, Meinhardt *et al.*, 2012). In general, the repressors showed complicated, multi-phasic kinetics which could not be used to determine quantitative association and dissociation rate constants. However, when the protein concentrations were fixed at low values, the amplitudes at equilibrium correlated with the percent of bound repressor predicted from filter binding assays (Figure 7 and Supplementary Figures 5 and 6). Therefore, changes in amplitude were used to compare differences in DNA binding affinity that resulted from (i) varied operator sequences or (ii) the presence of small molecules.

Other control experiments assessed the extent of nonspecific repressor binding to the streptavidin tips in the absence of immobilized DNA. A variety of measurements in the presence and absence of BSA indicated that the repressors probably had more nonspecific binding to the tips in the absence than in the presence of DNA. Since the amplitude of LacI binding to various DNA operators in the presence and absence of small molecule inducer matched all expectations, we conclude that background binding does not interfere with the assay when the protein concentration is held constant.

The performance of biolayer interferometry experiments could also be improved by pretreatment of the Cra and FruK proteins. For Cra, the amplitude of the biolayer interferometry signal was enhanced by exchanging the KP purification buffer for the PYK enzyme assay buffer using VivaSpin concentrators with a MWCO of 10,000 Da. To that end, the Cra protein sample was diluted 1:5 with buffer and concentrated ∼15 fold; this was repeated two more times prior to experiment. Since the final Cra concentration in the binding experiment was unchanged, we hypothesize that one of the KP buffer components diminished DNA binding activity; similar buffer effects were seen in the filter binding assay.

For FruK, aliquots that eluted at the leading edge of the peak from the DEAE column had some contaminant that was not present in the lagging edge. In biolayer interferometry experiments, this contaminant appeared to stimulate catalysis of F-1,6-BP to create the strong inducer F-1-P (Supplementary Figure 10). The contaminant was separated from the FruK using a VivaSpin concentrator with a MWCO of 10,000 Da to carry out buffer exchange into fresh FruK purification buffer, using the same dilution parameters as Cra buffer exchange. The resulting FruK had the same behavior towards FBP as protein purified from lagging end of the peak (Supplementary Figure 10). Adding the filtrate back to the reaction reproduced the apparent stimulation of catalysis for F-1,6-BP. These observations lead us to hypothesize that the affected FruK co-purifies with bound ADP (the other required substrate for the reverse catalytic reaction), which would indicate very high affinity of FruK for ADP.

For the Cra and Cra/FruK measurements, a day’s experiments typically used one operator DNA binding to: Cra, Cra+F-1-P, Cra+F-1,6-BP, Cra+FruK, Cra+FruK+F-1-P, and Cra+FruK+F-1,6-BP. Each set of experiments was repeated on separate days with different Cra and FruK preparations. From these, we determined that the day-to-day reproducibility of the equilibrium amplitude was ∼20% and that trends observed within each day’s work were reproducible on separate days. Repetitions used either of two lengths of DNA, which had the operator sequence embedded in either a 45mer or 69mer sequence. No DNA length dependence was observed.

For most experiments, Cra concentration was 50 nM. When present, FruK concentration was 200 nM and generally constituted ∼ 1/15^th^ of the final reaction volume. Because the FruK storage buffer differed significantly from the Cra storage buffer and contained 20% glycerol, we performed control experiments comprising Cra (i) pre-incubated with equivalent volumes of FruK storage buffer and (ii) with added glycerol; no effects were observed on DNA binding.

When fructose metabolites were included in the Cra and Cra/FruK DNA binding reactions, their final concentrations were 1 mM F-1-P and 10 mM F-1,6-BP unless otherwise indicated. When the ATP and ADDP nucleotides were included in the reactions, final concentrations were 3 mM unless otherwise indicated.

In several experiments with the Cra/FruK complex, we noted that the signal at “equilibrium” reached a maximal value and then diminished. Repetitions with freshly thawed aliquots showed this was due, at least in part, to FruK inactivation and/or destabilization over time. Interestingly, Cra/FruK samples with F-1,6-BP showed much less propensity for FruK inactivation.

## Acknowledgements

This project was supported by a grant (1 P30 GM110761) from the National Institute of General Medical Sciences at the National Institutes of Health to the Center for Biomedical Research Excellence in Protein Structure and Function, by NIH GM079423 to LSK, and by private funds. We thank Drs. Antonio Artigues and Maite Villar (KUMC) for assistance with mass spectrometry. We thank Drs. Maria Veiga-da-Cunha and Emile Van Schaftingen (Université Catholique de Louvain, Belgium) for the gift of the FruK expression plasmid. We thank Ms. Brittany Arce for assistance with growth assays and Drs. John Karanicolas and Lynn Hancock (KU-Lawrence) for the use of their plate readers. We thank Drs. Mark Fisher, Subhashchandra Naik, Joe Lutkenhaus, and Shishen Du (all of KUMC) for assistance with the biolayer interferometry assay and the use of their BLItz instruments. We thank Dr. Philip Gao and Ann Cooper (Protein Purification Core, KU-Lawrence, COBRE in Protein Structure and Function) for assistance with the heparin purification column. Dr. Alexey Ladokhin (KUMC) provided invaluable assistance translating Molchanova *et al.* (Molchanova *et al.*, 1992) from Russian to English. We thank Drs. Philip Hardwidge (Kansas State University), Sarah Bondos (Texas A&M Health science Center) and Jeffrey Bose (KUMC) for many helpful and interesting discussions about this work.

### Author Contributions

D.S. optimized the FruK and Cra protein purifications, carried out the FruK enzymatic assays, and assisted with manuscript development. MSF developed and carried out the biolayer interferometry assays. KSH carried out growth assays. CW helped to develop the Cra purification and performed the DNA binding assays using the filter binding technique. SM identified the LLhF-FruK and Cra-FruK interactions. ST performed the early LLhF/Cra purification and DNA binding assays. BFR prepared the F-1-P and developed the enzyme assay. PSH supervised growth assays and contributed to the manuscript. AWF designed and supervised enzyme assays and contributed to the manuscript. LSK supervised the project, participated in biolayer interferometry assays, performed data analyses, and wrote the manuscript.

## Footnotes

a They also noted that the *fruK* mutation rendered *E. coli* sensitive to mannose and xylitol toxicity.

